# Multi-omics profiling reveals microenvironmental remodeling as a key driver of house dust mite-induced lung cancer progression

**DOI:** 10.1101/2025.09.24.678347

**Authors:** Shams Al-Azzam, Isabella Stuewe, Sunandini Sharma, Miki Yamada-Hara, Arisachi Tanaka, Kegan Stringer, Merna Behnam, Norah Al-Azzam, Shuvro Nandi, Maria Zhivagui, Janelle Duong, Ting Yang, Scott Herdman, Maripat Corr, Nicholas J.G Webster, Eyal Raz, Ludmil B Alexandrov, Samuel Bertin

**Author notes:** Correspondence: Samuel Bertin, Ludmil Alexandrov.

## Abstract

Chronic exposure to the common aeroallergen house dust mite (HDM) induces lung inflammation and DNA damage, but its impact on lung cancer development remains largely unexplored. Using whole-genome sequencing, RNA-seq, and DNA methylation profiling, we assessed HDM effects in lung epithelial cell lines and a mouse orthotopic lung cancer model. HDM accelerated tumor growth without altering mutational burden. Transcriptomic and epigenetic analyses revealed tissue-specific effects: in normal lung, HDM enhanced pro-inflammatory and immune activation programs, whereas in tumors it suppressed T cell responses, antigen presentation, and chemokine signaling. Immune deconvolution showed a shift toward myeloid enrichment and lymphoid suppression, with reduced cytotoxic T and NK signatures. Notably, HDM-driven tumor promotion was abolished in *Il17a^−/−^* but not *Il1b^−/−^*mice, identifying IL-17A as a critical mediator. These findings demonstrate that chronic aeroallergen exposure reshapes the lung microenvironment to promote immune suppression and accelerate lung cancer progression.

**Highlights:** Chronic house dust mite (HDM) exposure accelerates lung tumor growth through non-mutagenic, immune-mediated mechanisms.

HDM activates pro-inflammatory and immune programs in normal lung tissue but suppresses antitumor T cell responses in tumors.

Multi-omics profiling reveals epigenetic silencing of immune genes and a myeloid-enriched, lymphoid-deficient tumor microenvironment.

HDM-driven tumor promotion depends on IL-17A but not IL-1β, establishing IL-17A as a central driver of lung tumor promotion.

## Introduction

House dust mite (HDM) is one of the most prevalent indoor aeroallergens and has been implicated in the pathogenesis of asthma^1^. Exposure to allergens from HDM species such as *Dermatophagoides pteronyssinus* elicits T helper 2 (Th2) immune responses, characterized by eosinophilic infiltration and mucus overproduction^1^. Chronic HDM exposure also promotes Th17 responses, neutrophilic infiltration, and airway remodeling^1,2^. In addition to its pro-inflammatory effects, HDM induces oxidative stress and DNA double-strand breaks in airway epithelial cells, leading to DNA damage^3,4^.

Chronic inflammation is increasingly recognized as a contributing factor in cancer initiation and progression, particularly in the lung^5^. Inflammatory mediators can influence the tumor microenvironment (TME), alter immune surveillance, and drive genomic instability^5^. However, the contribution of HDM-induced inflammation and DNA damage to lung tumorigenesis remains underexplored, and it is largely unknown whether HDM exposure drives mutagenesis, epigenetic remodeling, or alterations in the TME.

Most studies to date have focused on HDM’s immunologic effects in asthma, leaving a critical knowledge gap regarding its role in lung tumor biology. Determining whether HDM exposure alters the genomic or epigenomic landscape of normal lung epithelial and tumor cells is essential to uncover mechanisms by which chronic aeroallergen exposure may promote lung carcinogenesis.

Here, we evaluate the impact of HDM exposure on the genomic, epigenomic, and transcriptomic landscapes of normal lung epithelial and tumor cells. HDM effects were assessed *in vitro* using lung epithelial cell lines and *in vivo* in a mouse orthotopic lung tumor model. We analyzed mutational burden and signatures across both systems and performed integrated epigenomic, transcriptomic, and immune profiling of tumors and matched normal lung tissues to comprehensively characterize the effects of HDM exposure.

## Results

### Chronic HDM exposure accelerates lung carcinoma progression

To evaluate the potential tumor-promoting effects of chronic HDM exposure, we employed a syngeneic orthotopic lung cancer model using Lewis lung carcinoma (LLC) cells^6^. Wild type (WT) C57BL/6J mice were sensitized by intranasal (i.n.) instillations with HDM or vehicle (VEH) once on Day 0, followed by three i.n. challenges per week for four weeks. On Day 36, 24 hours after the final i.n. challenge, mice received orthotopic lung tumor implantation of luciferase-expressing LLC (LLC-GLF) cells via i.n. administration. Tumor growth was monitored by bioluminescence imaging (BLI) using an *in vivo* imaging system (IVIS), as previously described^7^. On Day 64, 24 hours after the last BLI, mice were euthanized, and lungs were harvested for analysis **(Figure 1A)**. Mice chronically exposed to HDM exhibited significantly higher tumor bioluminescence signals between 3 and 4 weeks after LLC cell implantation compared to VEH-treated controls **(Figures 1B, 1C, and S1)**. At endpoint, gross examination revealed macroscopic lung tumors in both VEH- and HDM-treated mice. However, HDM-treated mice showed more extensive disease, with visibly enlarged lungs **(Figure 1D)** and significantly increased lung weight **(Figure 1E)**. To further assess tumor burden, we quantified lung tumors on scanned hematoxylin and eosin (H&E)-stained sections using QuPath software^8^. In line with the increased bioluminescence signal and lung weight, HDM-treated mice exhibited greater tumor multiplicity (defined as the total number of lung tumors per mouse) and tumor area (defined as the cumulative surface area of all tumors) compared with VEH-treated controls **(Figures 1F–H)**.

**Figure 1.**
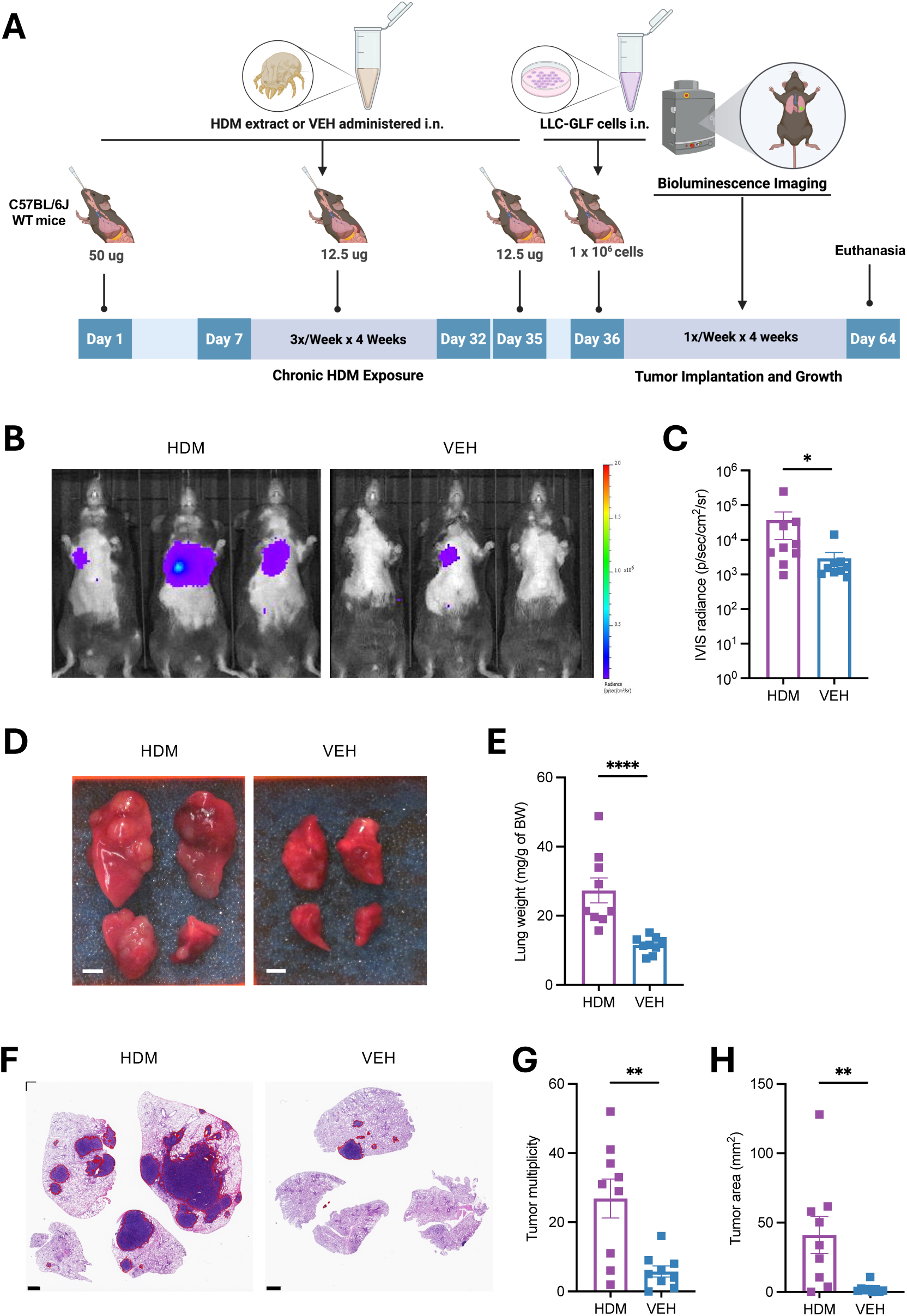
Chronic Exposure to HDM Accelerates Tumor Growth in a Mouse Model of Orthotopic Lung Cancer. **A)** Age-matched male C57BL/6J WT mice (n = 9 mice/group) were first sensitized by intranasal (i.n.) instillations under light isoflurane anesthesia with HDM (50 µg/50 µL per mouse) or vehicle control (VEH; 50 µL per mouse) on Day 0, followed by three i.n. challenges per week (M/W/F) for four weeks with HDM (12.5 µg/50 µL per mouse) or VEH (50 µL/mouse). LLC-GLF cells (1 × 10^6^) were administrated i.n. on Day 36, 24h after the last HDM or VEH i.n. challenge, and tumor growth was followed by bioluminescence imaging (BLI) using an *in vivo* imaging system (IVIS), as outlined in the schematic overview of the study design. On Day 64, 24h after the last BLI the mice were euthanized, and lungs were harvested for analysis. **B)** Representative IVIS images of tumor signals on Day 63 from three representative mice per group. Tumor growth kinetics for all experimental mice are shown in Figure S1. **C)** IVIS radiance at Day 63 in mice treated with either HDM (purple) or VEH (blue). **D)** Representative photos of 4 lung lobes (dorsal view). Scale bars, 0.25 cm. **E)** Lung weight normalized to mouse body weight (BW). **F)** Representative photos of H&E-stained lung sections in the two experimental groups. The areas within the red borders were considered positive for tumors. Scale bars, 1 mm. **G)** Tumor multiplicity (the total number of lung tumors per mouse) calculated on H&E-stained sections as shown in F. **H)**Tumor area (the cumulative surface area of all tumor per mouse) calculated on H&E-stained sections as shown in F. Data are presented as mean ± SEM. Statistical significance was assessed by Mann-Whitney test; * *p* <0.05, ** *p* <0.01, **** *p* <0.0001.

### HDM does not have mutagenic effects in lung epithelial cell lines *in vitro*

HDM has been reported to induce double-stranded DNA breaks in airway epithelial cells^3,4^, a genotoxic effect that, if not properly repaired, can result in permanent somatic mutations and potentially contribute to tumor initiation and progression. To evaluate the genotoxic and mutagenic potential of HDM, we first examined its effects *in vitro* using human and mouse lung epithelial cell lines. BEAS-2B human lung epithelial cells and LLC mouse lung carcinoma cells were treated with increasing concentrations of HDM for 24 hours, and cell viability was assessed. HDM reduced cell viability in a concentration-dependent manner, with half-maximal inhibitory concentrations (IC₅₀) of ∼200 µg/mL in BEAS-2B cells and ∼400 µg/mL in LLC cells **(Figure 2A)**, consistent with previous reports^3,4^. Cells were then treated with HDM (at their respective IC_50_ concentration) or vehicle (PBS) for 24 hours, expanded, and subjected to whole-genome single-molecule duplex sequencing **(Figure 2B)**. This approach enables detection of true single-molecule mutations without clonal expansion, providing a direct and highly accurate measure of mutagenesis^9^. In both cell lines, an average of one billion duplex molecules was sequenced per sample; however, mutational burden did not differ between HDM-and PBS-treated cells **(Figure 2C)**.

**Figure 2.**
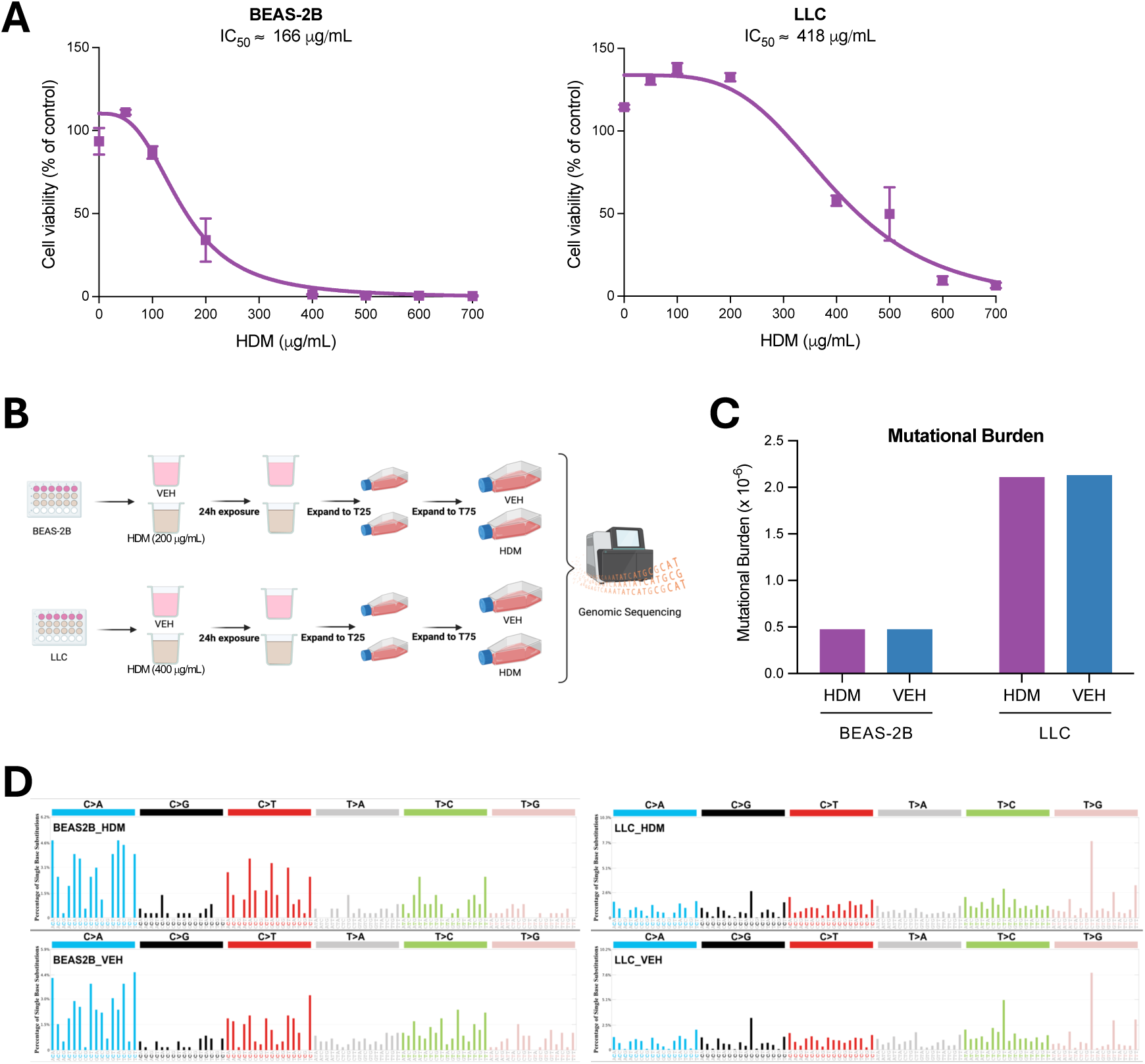
HDM Does Not Have Mutagenic Effects in Human and Mouse Lung Epithelial Cell Lines *In Vitro*. **A)** Dose-response curves of HDM treatment in BEAS-2B and LLC cells. Cell viability is shown relative to vehicle control (PBS; % of control). Calculated IC_50_ values are indicated. **B)** Experimental setup for HDM exposure and duplex sequencing. BEAS-2B and LLC cells were treated with either HDM (200 µg/mL for BEAS-2B; 400 µg/mL for LLC) or vehicle (VEH; PBS), expanded across multiple passages, and subsequently subjected to genomic sequencing. **C)** Comparison of mutational burden between samples treated with either HDM (purple) or VEH (blue), shown as somatic mutations per megabase (Mb) of coding DNA. **D)** Mutational spectra of BEAS-2B and LLC cells treated with HDM compared to VEH controls. Mutations are classified into the six base substitution categories and displayed in SBS96 format. The experiments were performed in triplicate (A) or singlicate (B-D).

To further characterize mutational patterns, we generated single base substitution (SBS) profiles using the SBS96 classification schema, which categorizes mutations into 96 classes defined by substitution type (C>A, C>G, C>T, T>A, T>C, and T>G) and flanking sequence context^10,11^. We also analyzed double base substitutions (DBSs), which represent simultaneous substitutions of two adjacent nucleotides and are classified into 78 categories by dinucleotide context^10,11^ and small insertions and deletions (indels), which are grouped into 83 categories^10,11^. Both DBS and indel profiles were similar between HDM- and PBS-treated cells **(Figure S2)**. Comparative analysis revealed highly concordant mutational patterns (cosine similarity > 0.98) between treatment groups in both BEAS-2B and LLC cells, suggesting that direct HDM exposure does not significantly alter mutational processes under these experimental conditions **(Figure 2D)**.

### HDM exposure does not increase mutagenesis in mice *in vivo*

We next assessed whether chronic HDM exposure promotes mutagenesis in the LLC orthotopic lung cancer model by treating an independent cohort of mice as in **Figure 1A**. At endpoint, macroscopic tumors and normal adjacent lung tissues were harvested for analysis: tumor tissues were subjected to bulk whole-genome sequencing (WGS; 90x coverage), while normal lung tissues were analyzed using single-molecule duplex sequencing, with an average of one billion duplex molecule profiled per sample **(Figure 3A)**. Genomic DNA from each mouse’s spleen was used as a germline reference (30x WGS), and common variants across mice were removed to generate an HDM-specific profile. Consistent with the *in vitro* findings, quantification of mutational burden revealed no statistically significant differences between HDM- and VEH-treated groups in either normal lung tissues or tumors **(Figure 3B)**. Although a trend toward higher mutational burden was observed in HDM-treated samples, the differences did not reach statistical significance (*p* = 0.31 for both tissues), possibly due to limited sample size (n = 3) or reflecting only a weak mutagenic effect. Mutational spectrum analysis supported these observations. SBS96 profiles were highly similar between HDM- and VEH-treated samples (cosine similarity > 0.98; **Figures 3C and 3D)**, and the distribution of mutation types and trinucleotide contexts remained unchanged. Together, these results suggest that chronic HDM exposure does not significantly increase mutational burden or alter mutational processes in either the lung microenvironment or tumor cells under our experimental conditions.

**Figure 3.**
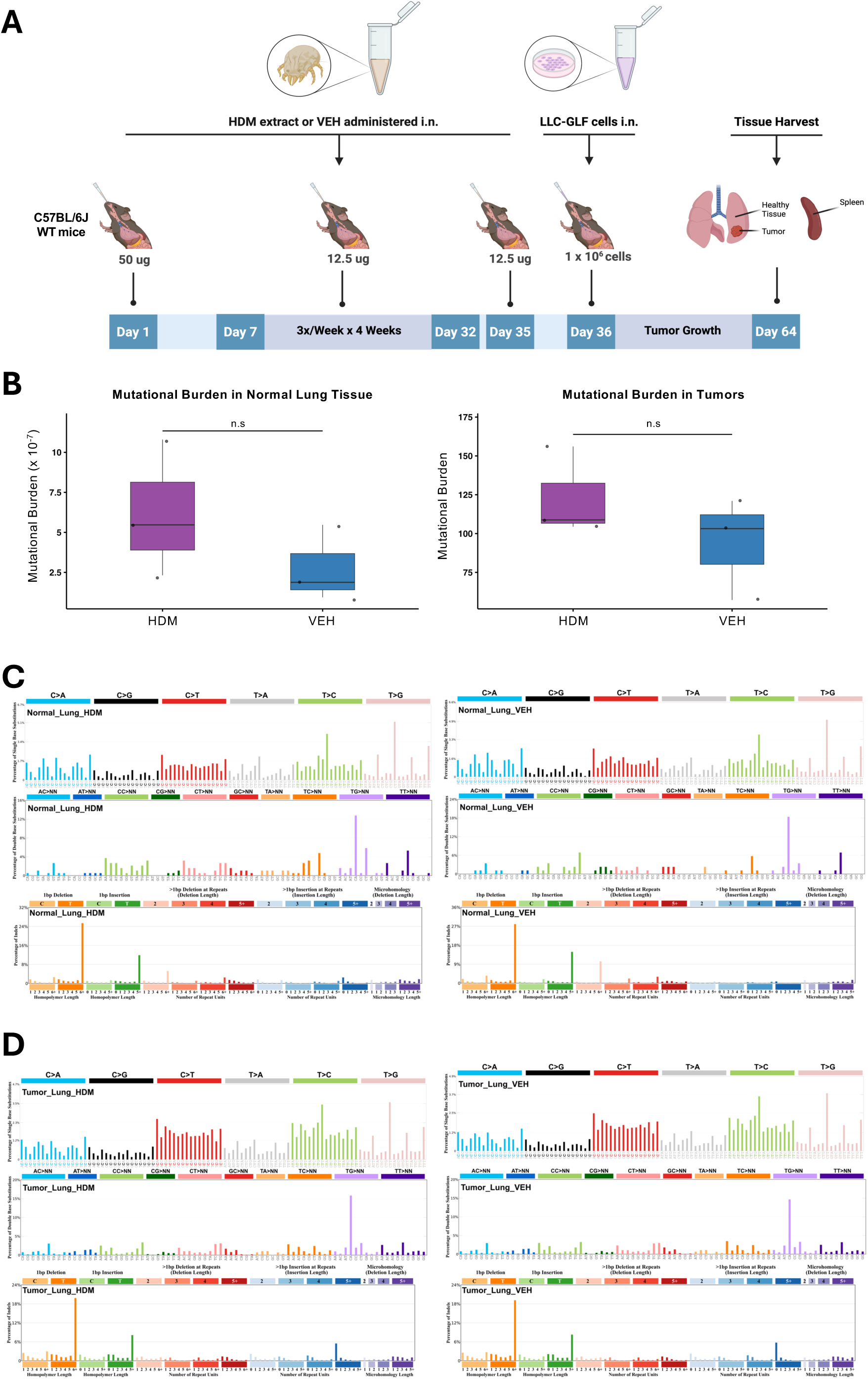
HDM Does Not Have Mutagenic Effects in Normal Lung Tissues or Tumors in Mice *In Vivo*. **A)** C57BL/6J WT mice (n = 3 mice/group) were first sensitized by i.n. instillations under light isoflurane anesthesia with HDM (50 µg/50 µL per mouse) or vehicle control (VEH; 50 µL per mouse) on Day 0, followed by three i.n. challenges per week (M/W/F) for four weeks with HDM (12.5 µg/50 µL per mouse) or VEH (50 µL/mouse). LLC-GLF cells (1 × 10^6^) were administrated i.n. on Day 36, 24h after the last HDM or VEH i.n. challenge. Mice were euthanized on Day 64, and the lungs and spleens were harvested for analysis as outlined in the schematic overview of the study design. **B)** Mutational burden in normal lung tissues (left) and tumors (right) from mice treated with either HDM (purple) or VEH (blue). Mutational burden is expressed as mutations per megabase (Mb). Boxes show the median (center line) and interquartile range (IQR); whiskers extend to 1.5× the IQR; points are individual samples (HDM, n = 3; VEH, n = 3). *p-*values were computed using a two-sided Welch’s *t*-test. n.s = not significant. **C)** Mutational spectra in normal lung tissues. Single base substitutions (SBS96), insertion-deletions (ID83), and doublet base substitutions (DBS78) are shown for HDM- and VEH-treated samples. **D)** Mutational spectra in tumor tissues. SBS96, ID83, and DBS78 profiles are shown for HDM- and VEH-treated samples. The average profiles of three tumors from independent mice are shown in panels C and D.

### HDM exposure alters gene expression in tumors and normal lung tissues

Because the tumor-promoting effect of HDM exposure could also be driven by non-mutagenic mechanisms, we next evaluated the transcriptomes of tumors and adjacent normal lung tissues using bulk RNA sequencing (RNA-seq). As shown in the principal component analysis (PCA) **(Figure S3A)**, normal lung tissues are distinctly separated between HDM and VEH treated. In contrast, tumor tissues exhibit partial cluster overlap between HDM and VEH, suggesting that HDM-induced transcriptional changes are more pronounced in normal lung tissues than in tumors, where gene expression programs may already be dysregulated. Samples were clustered by hierarchical clustering based on log2(transcripts per million [TPM]+1) **(Figures S3B and S3C)**. As shown in the volcano plots **(Figure 4A)**, HDM-treated normal lung tissue exhibited 1,390 differentially expressed genes (DEGs) compared to VEH-treated controls, with 1,004 genes upregulated and 386 genes downregulated (absolute fold change ≥ 2, false discovery rate [FDR] < 0.05). In tumor tissue treated with HDM compared to VEH controls, 599 DEGs were identified, including 449 upregulated and 150 downregulated DEGs. These findings suggest that, in our experimental model, the normal lung microenvironment is more transcriptionally responsive to HDM than tumor tissue.

**Figure 4.**
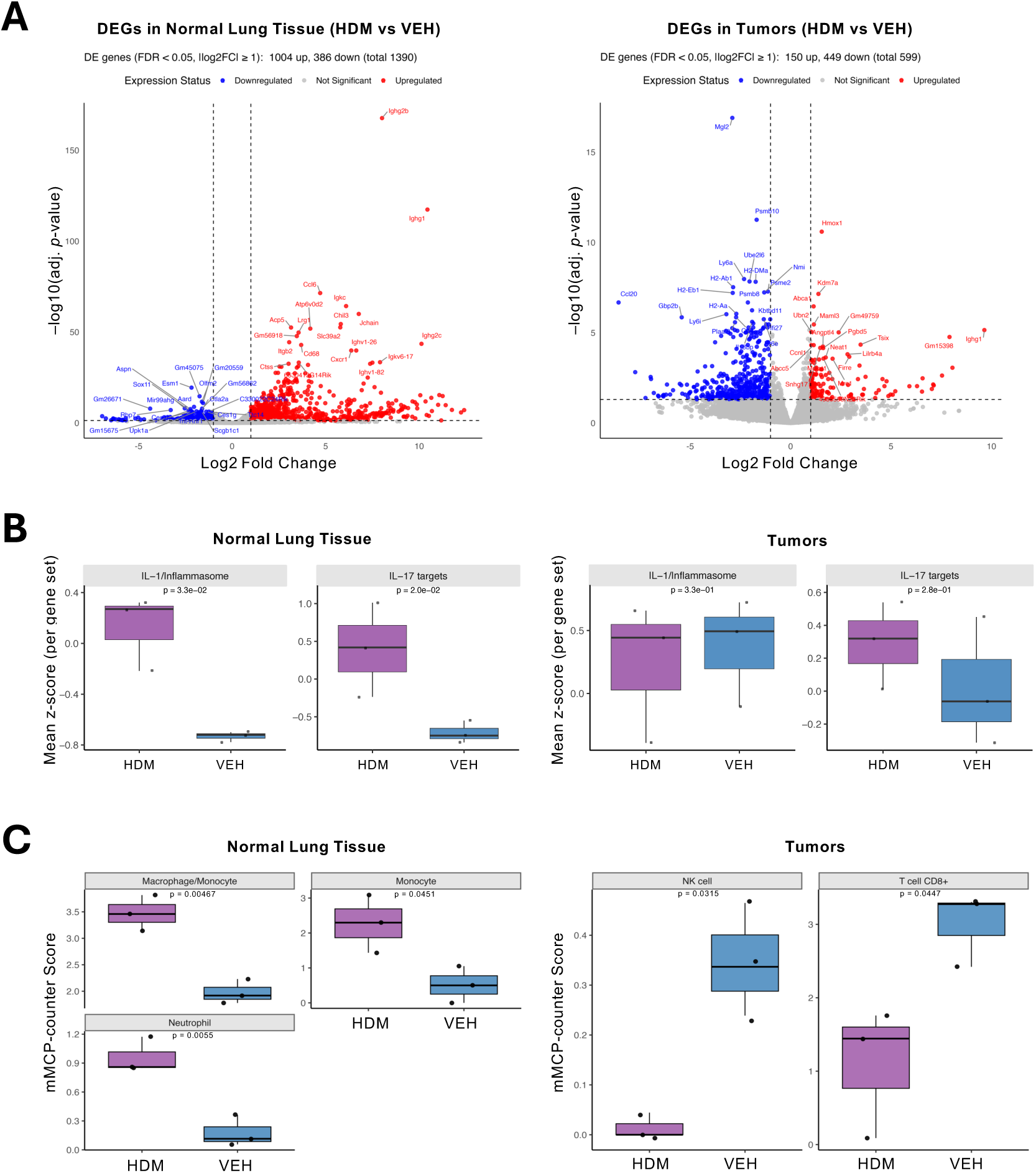
HDM Exposure Differentially Modulates Gene Expression and Immune Cell Programs in Tumors and Normal Adjacent Lung Tissues. **A)** Differentially expressed genes (DEGs) in normal (left) and tumor (right) lung tissue comparing HDM-versus VEH-treated samples. Volcano plots display significantly upregulated (red) and downregulated (blue) genes (FDR < 0.05, |log2 fold change| ≥ 1). **B)** Boxplots summarize mean z-scores (per gene set) for curated IL-1/inflammasome- and IL-17-related target genes in normal lung tissues (left panels) and tumors (right panels) from mice treated with either HDM (purple) or VEH (blue). Higher z-scores reflect increased module signature activity. Each dot represents an individual sample. **C)** Boxplots depict mMCP-counter scores for the indicated significant (*p* < 0.05) cell populations in tumors (left) and normal lung tissues (right) from mice treated with either HDM (purple) or VEH (blue). Boxes show the median (center line) and interquartile range (IQR); whiskers extend to 1.5× the IQR; points are individual samples (HDM, n = 3; VEH, n = 3). *p-*values were computed using a two-sided Welch’s *t*-test.

As shown in the Venn diagrams **(Figure S3D),** analysis of gene expression in HDM-treated samples identified 32 DEGs commonly upregulated in both normal lung tissues and tumors. These included key inflammatory and immune response genes such as *Hmox1*, *Il33*, *Cxcl1*, *Mrc1*, and *Saa3* **(Figures 4A** **and S3E)**. Several immunoglobulin-related genes (*Igkv*, *Ighv*, *Ighg*) were also upregulated, suggesting activation of local B cell responses. The shared upregulation of these genes indicates that HDM exposure triggers a conserved immune and epithelial stress response in both normal lung tissues and tumors. We also identified a shared set of 25 genes that were significantly downregulated in both normal lung tissues and tumors following HDM exposure **(Figure S3D)**. These included transcriptional regulators (*Gata3, Smad6, Ripply3*), mitochondrial genes (*mt-Rnr1, mt-Rnr2, mt-Nd2-4*), and structural or epithelial-associated genes (*Prom2, Egfl7, Tmem47, Tnxb)* **(Figure S3E)**. These findings suggest that HDM exposure reprograms the transcriptome by simultaneously enhancing inflammatory pathways and dampening epithelial homeostasis, mitochondrial activity, and developmental transcriptional programs across both tissue types.

### HDM exposure induces distinct interleukin gene signatures in normal lung tissues and tumors

To evaluate how HDM exposure impacts cytokine signaling in the lung, we examined expression of genes encoding cytokines and cytokine receptors in both normal lung tissues and tumors. In normal lung tissues, HDM exposure led to significant upregulation of several interleukin (IL)-related genes, including *Il1b, Il1r2*, *Il2rb*, *Il16*, *Il17ra*, *Il18*, *Il18rap*, and *Il33* **(Figure S4A)**. These genes are associated with pro-inflammatory signaling and Th2/Th17 immune responses, indicating that HDM primes the lung environment through immune activation and cytokine receptor upregulation.

In lung tumors, *Il1rap* and *Il33* were also significantly upregulated upon HDM exposure, together with *Il1r1*, while *Il18bp*, *Il1b*, *Il2rb*, and *Il4i1* were significantly downregulated compared to tumors from VEH-treated mice **(Figure S4B)**. The distinct cytokine profile in tumor versus normal tissues in response to HDM indicates that HDM exposure differently affects each tissue. Notably, the decreased expression of *Il1b*, a major pro-inflammatory cytokine, and *Il18bp*, a decoy receptor that neutralizes IL-18 activity, in HDM tumors suggests a reprogramming of immune responses within the TME toward a more immunosuppressive state.

Since several of the modulated cytokines are products of inflammasome activation, we next assessed the expression of inflammasome-related genes. While tumors showed no significant changes, *Aim2*, *Nlrp3*, *Nlrp12*, and *Casp1* were significantly upregulated in normal lung tissue following HDM exposure **(Figure S4C)**, consistent with the increased expression of *Il1b*, *Il18*, and *Il33* in these samples.

In line with these findings, IL-1/inflammasome- and IL-17-related target gene module scores were significantly higher in HDM-exposed normal lung tissue compared with VEH **(Figure 4B)**. Gene set enrichment analysis (fgsea) confirmed significant upregulation of both an IL-1/inflammasome module (NES = 1.64, FDR = 0.026) and an IL-17 module (NES = 1.65, FDR = 0.026) in normal lung tissue. Module scoring indicated that the IL-17 signal was driven by canonical effector chemokines (*Cxcl1*, *Cxcl2*, *Lcn2*, *S100a8/9*), whereas IL-1 enrichment reflected inflammasome machinery (*Casp1*, *Nlrp3*, *Il1r1*, *Il1b*), consistent with robust HDM-induced activation of both pathways in normal lung tissue. In contrast, tumors displayed constitutively elevated IL-1/inflammasome scores irrespective of treatment, while IL-17 module scores increased only modestly with HDM exposure. Neither IL-17 nor IL-1 modules were significantly enriched in tumors (IL-17 NES = -1.02, FDR = 0.45; IL-1 NES = 1.14, FDR = 0.45), and module scores were comparable between HDM and VEH groups. These results suggest that IL-1 and IL-17 programs in tumors are largely constitutively activated or restricted to the microenvironment, with HDM driving only a limited IL-17-associated response.

### HDM exposure enhances T cell responses in normal lung tissues but suppressed them in tumors

To assess how HDM exposure influences adaptive immune responses in tumor-bearing lungs, we compared normal lung tissues and tumors in HDM- and VEH-treated mice, focusing on curated Th1-, Th2-, and Th17-associated cytokines, chemokines, transcription factors, and TNF/TNFR family members **(Figure S5)**. HDM exposure substantially enhanced adaptive T cell responses in normal lung tissue, with broad upregulation of Th1-, Th2-, and Th17-associated genes. Within the Th1 program, multiple TNF superfamily co-stimulatory molecules (*Tnfsf9, Tnfrsf1b, Tnfrsf14, Tnfrsf26*) and select chemokines (*Cxcl9, Ccl5*) were upregulated, along with transcription factors essential for Th1 differentiation (*Tbx21, Stat1*). Th2 responses were partially activated, evidenced by upregulation of TNF/TNFR members (*Tnfrsf25, Tnfrsf13b*), whereas the canonical Th2 marker *Gata3* was downregulated. For the Th17 axis, HDM induced strong upregulation of chemokines (*Ccl20*), key transcriptional drivers for Th17 lineage (*Stat3*, *Batf*), TNF superfamily members (*Tnfrsf9*, *Tnfsf13b*), and *Nlrp3*, which contributes to Th17 stability^12,13^. In contrast, tumors from HDM-treated mice displayed broad suppression of Th1 and Th2 responses. Key Th1-associated chemokines (*Cxcl9*, *Ccl5*), transcription factors (*Tbx21*, *Stat1*, *Irf1*, *Irf8*), and TNF superfamily members (*Tnfsf10*, *Tnfrsf9*) were significantly downregulated, while Th2-related chemokines (*Ccl17*, *Ccl22*) and the co-stimulatory receptor *Tnfrsf4* were also suppressed. The Th17 response appeared attenuated, with *Ccl20* and *Batf* downregulated, whereas *Stat3*, *Tnfsf13b*, and *Nlrp3* showed no positive log2 fold change. Collectively, these findings suggest that HDM enhances T cell responses in normal lung tissue, whereas such responses are actively suppressed within tumors.

### HDM exposure reshapes the TME

Deconvolution using the murine Microenvironment Cell Populations-counter (mMCP-counter), a tool to estimate the immune and stromal composition of heterogeneous tissue from transcriptomic data^14^, revealed distinct immune landscapes in normal lung tissues versus tumors. HDM-exposed normal lung tissues showed increased macrophage, monocyte, and neutrophil scores compared to VEH. In contrast, tumors from HDM-treated mice displayed significantly decreased natural killer (NK) cell and CD8^+^ T cell scores **(Figure 4C)**, suggesting impaired antitumor cytotoxic responses. Under HDM exposure, tumors also showed a significant reduction in B cell and neutrophil abundance relative to matched normal tissue, whereas macrophage/monocyte scores were elevated in tumors across both treatment groups **(Figure S6)**. In VEH controls, tumors displayed higher cancer-associated fibroblast (CAF), endothelial cell, and T cell scores compared with normal tissue. Collectively, these findings indicate a robust, exposure-independent myeloid enrichment in tumors, with HDM specifically suppressing B cell, CD8⁺ T cell, NK cell, and neutrophil compartments, thereby promoting a more immunosuppressive TME.

### HDM exposure drives divergent immune and metabolic pathways in normal lung tissues and tumors

To identify biological pathways affected by HDM exposure in lung tumors and normal adjacent tissues, we performed pathway enrichment analysis on DEGs using ConsensusPathDB^15^, KEGG^16^, Reactome^17^, and MouseCyc^18^ pathway databases. In normal lung tissues, HDM induced strong activation of innate and adaptive immune pathways, including Fc-gamma receptor (FCGR) signaling, neutrophil degranulation, and B cell receptor signaling **(Figure 5A)**. Cell surface interactions at the vascular wall, along with other pathways not included in the curated set, such as osteoclast differentiation and hemostasis, were also enriched **(Figure S7A)**, suggesting increased levels of inflammation and tissue remodeling. However, the pathway labeled "osteoclast differentiation" may reflect activation or differentiation of lung myeloid cells or tumor-associated macrophages (TAMs), as osteoclasts derive from the myeloid lineage and share many genes with macrophages and dendritic cells. Indeed, genes like *Trem2*, *Syk*, *Plcg2*, *Spi1* (PU.1), and Fc receptors (*Fcgr1*, *Fcgr2b*, *Fcgr3*, *Fcgr4*) are highly expressed in macrophage subsets involved in lung immune responses and tumor-associated inflammation. Additionally, the presence of *Il1b*, *Socs3*, *Ncf1/2/4* (components of the NADPH oxidase complex), and *Btk* indicates inflammatory signaling and reactive oxygen species (ROS) generation, which are hallmarks of inflamed tissues and tumors. These processes likely prime the TME and underscore the substantial impact of HDM exposure on normal lung tissue.

**Figure 5.**
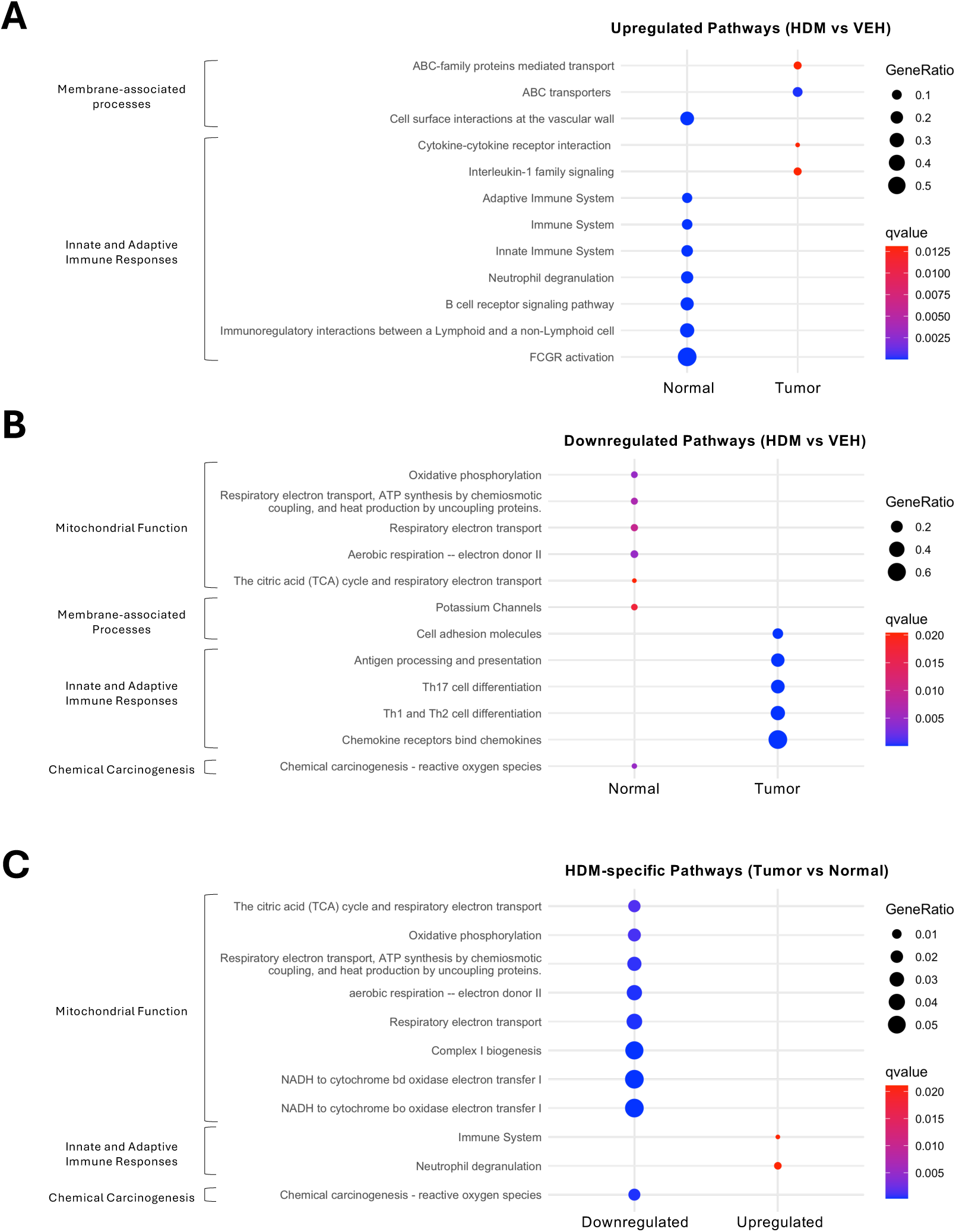
HDM Exposure Alters Signaling Pathways in Tumors and Normal Adjacent Lung Tissues. Pathway enrichment analysis of DEGs identified in tumor and normal adjacent lung tissues based on KEGG, Reactome, and MouseCyc pathway databases. The dot plots display curated pathways that are either **A)** upregulated or **B)** downregulated in normal lung tissues or tumors following HDM exposure compared to VEH. **C)** Common pathways modulated by HDM in both normal lung tissues and tumors. Dot size represents the number of DEGs associated with each pathway, while dot color indicates statistical significance (*q*-value, calculated using the Benjamini-Hochberg false discovery rate).

In contrast, tumors from HDM-exposed mice exhibited a more limited transcriptional response, with modest enrichment of cytokine-cytokine receptor interactions, IL-1 signaling, and growth hormone/circadian rhythm-related pathways **(Figures 5B** **and S7A)**, but lacking the broader immune activation signatures seen in normal tissues. The modest enrichment of the IL-1 pathway, despite no difference in IL-1/inflammasome module scores between HDM- and VEH-exposed tumors **(Figure 4B)**, likely reflects that pathway enrichment incorporates all IL-1-related genes, including those constitutively expressed or outside the canonical inflammasome machinery, whereas module scoring captures coordinated changes in core inflammasome components. More strikingly, HDM exposure led to a significant downregulation of T cell-related pathways in tumors, including Th1, Th2, and Th17 cell differentiation, antigen processing and presentation, and chemokine signaling **(Figures 5B** **and S7B)**. This suppression of T cell responses and chemokine gradients may promote immune evasion and favor a more tolerogenic TME.

We next performed pathway enrichment analysis on shared DEGs to identify pathways commonly affected by HDM exposure in both lung tumors and normal adjacent tissues. HDM exposure consistently increased immune system activation and neutrophil degranulation pathways in both tissue types **(Figure 5C)**. Conversely, HDM-exposed tumors and normal adjacent tissues displayed decreased oxidative phosphorylation (OXPHOS), respiratory electron transport, and ROS-related pathways, indicating altered mitochondrial respiration and a potential metabolic shift of tumor cells toward aerobic glycolysis (Warburg effect), a hallmark of cancer^19^. Interestingly, pathways typically associated with neurodegenerative diseases (e.g., Alzheimer’s disease, Amyotrophic lateral sclerosis) were also suppressed **(Figure S7C)**, likely reflecting mitochondrial dysfunction signatures. Metabolic and thermogenesis pathways (e.g., Thermogenesis, Retrograde Endocannabinoid Signaling, Diabetic Cardiomyopathy) were also commonly downregulated in lung tumor and normal adjacent tissues, but these changes may be secondary to metabolic shifts in the lung microenvironment under chronic HDM exposure.

Collectively, these findings suggest that chronic HDM exposure induces divergent immune and metabolic responses in lung tumors and normal adjacent tissues, shaping both the lung and tumor microenvironments in ways that may promote tumor progression.

### HDM induces widespread promoter hypermethylation in normal lung but has limited epigenetic effects in tumors

Given that epigenetic alterations can contribute to transcriptomic changes, we next analyzed the DNA methylation profiles of tumors and normal adjacent tissues to assess whether chronic HDM exposure induces epigenetic remodeling. CpG sites were examined along with four additional genomic feature sets: genome-wide tiling regions of 5,000 bp (n = 129,034), Ensembl genes (version 75; n = 24,298), promoter regions of Ensembl genes (version 75; n = 27,522), and CpG islands from the UCSC Genome Browser database^20^ (n = 12,331) for both tissue types under HDM or VEH treatment.

Quantitative analysis of differentially methylated sites (DMSs; |Δβ| ≥ 0.2, *p* < 0.05) revealed a marked reduction in the number of altered sites in tumors compared with normal tissues for all genomic features **(Figure S8)**. In normal lung tissue, HDM exposure induced widespread methylation changes in promoters, with x hypermethylated and y hypomethylated sites relative to VEH. In contrast, tumors displayed markedly fewer promoter changes, with only x hypermethylated and z hypomethylated sites after HDM treatment. Density plots of β-values further revealed a clear shift in promoter methylation profiles in normal tissue following HDM exposure, whereas tumor promoter remained largely unchanged. Across other genomic features, normal tissue consistently exhibited more DMSs than tumor tissue, with a predominant hypermethylation bias in promoters, CpG islands, and tiling regions. Tumor tissue showed fewer changes overall and a more balanced hyper-to hypomethylation ratio. These results indicate that HDM exposure elicits a robust epigenetic response in normal lung tissue, particularly at promoter regions, whereas promoter methylation changes in tumors are comparatively limited. This reduced response may reflect pre-existing epigenetic alterations already established during tumorigenesis, such as global hypomethylation, localized promoter hypermethylation of tumor suppressor genes, or widespread chromatin remodeling that limit further epigenetic plasticity. Alternatively, the reduced response in tumors may be influenced by the timing of HDM exposure relative to tumor development, or by selection pressures within the TME that stabilize existing epigenetic programs.

### Epigenetic control of immune gene expression is preserved in normal lung but lost in tumors

Promoter methylation is a key mechanism for transcriptional regulation and tumor-suppressor gene silencing. To investigate the functional impact of HDM-induced methylation changes, we integrated promoter DNA methylation and gene expression in two steps. First, we quantified differentially methylated genes (DMGs) and DEGs independently, without requiring concordance between the two datasets. Genes were classified into four categories based on the intersection between the DMGs and DEGs: hypermethylated-upregulated (hyper-up), hypermethylated-downregulated (hyper-down), hypomethylated-upregulated (hypo-up), and hypomethylated-downregulated (hypo-down) genes **(Figure 6A)**. In normal tissue, promoter methylation and gene expression were strongly anti-correlated, whereas in tumor tissue, this relationship was weaker, showing a partial decoupling. Tumors were biased toward hypermethylation-mediated repression but also showed a greater number of discordant cases, suggesting that other regulatory mechanisms may predominate in the tumor context. Second, we filtered for genes showing inverse correlation between methylation and expression (i.e., hypermethylation with downregulation or hypomethylation with upregulation) to identify potential “epigenetically controlled” genes. In normal tissue, 20 genes met this criterion, including 12 epigenetically activated (hypomethylated/upregulated: *Ccl9*, *Trem2*, *Oas3*, *Tmem106a*, *Slfn8*, *Trem3*, *Lrrc25*, *Tarm1*, *Capg*, *Fcer1g*, *Osm*, *Bin2*) and 8 epigenetically repressed genes (hypermethylated/downregulated: *Gata3*, *Ripply3*, *Lrat*, *Pdlim3*, *Tagln*, *Tril*, *Lepr*, *Tubb4a*) **(Figure 6B)**. In contrast, no genes in tumor tissue passed this inverse-correlation filter, suggesting a loss of direct epigenetic control in tumors or the predominance of pre-existing silencing mechanisms.

**Figure 6.**
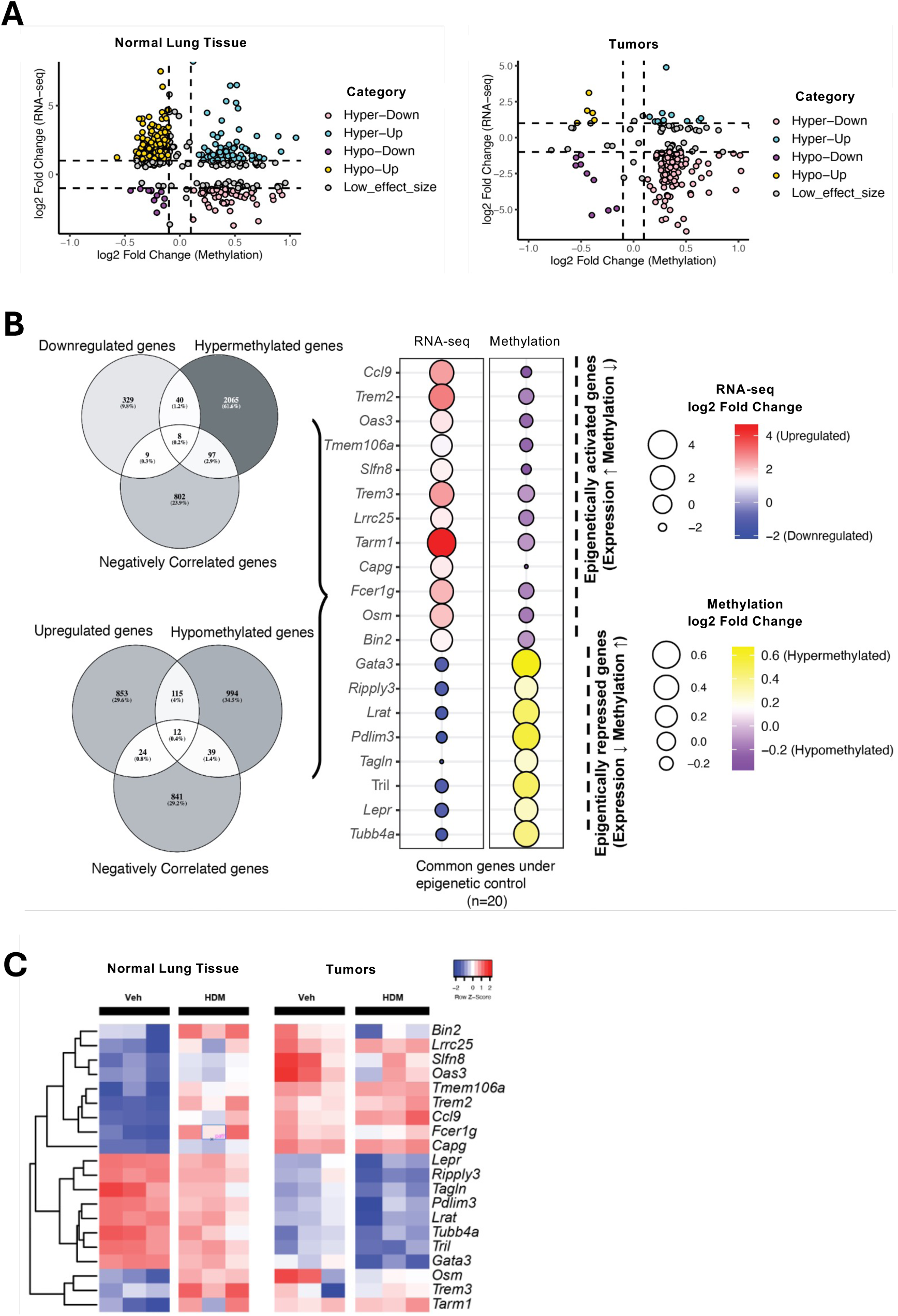
HDM Exposure Alters DNA Methylation Profiles in Normal Lung Tissues and Tumors. **A)** Integrated methylation-expression scatterplots. For each gene, log2 fold change in RNA-seq (y-axis) is plotted against log2 fold change in promoter methylation (x-axis; M-value scale). Genes are classified into four quadrants: hyper-down (hypermethylated & downregulated), hyper-up, hypo-down, and hypo-up. Dashed lines indicate calling thresholds for expression and methylation effect sizes and FDR as described in the corresponding Methods section. **B)** Genes under putative epigenetic control in normal tissue. Left, Venn diagrams show the overlap between differentially methylated genes (DMGs) and differentially expressed genes (DEGs). Right, dot plots display per-gene changes in RNA-seq (log2 fold change) and promoter methylation (log2 fold change, M-values) for the 20 inversely regulated genes in normal tissues; color encodes direction and magnitude (see scales). **C)** Expression heatmap for the 20 epigenetically regulated genes. Z-scored expression across samples is shown for normal and tumor tissues from mice treated with either VEH or HDM. Columns are grouped by tissue and exposure; genes are clustered by hierarchical clustering. DEGs and DMGs were called independently and then intersected; multiple testing was controlled by Benjamini-Hochberg false discovery rate (FDR < 0.05 unless noted). Effect sizes are log2 fold change for RNA-seq and log2 fold change of promoter methylation (M-values). Exact thresholds and promoter definitions are provided in Methods

Next, we tracked the expression and methylation changes of these 20 epigenetically regulated genes after VEH and HDM treatments in both normal and tumor tissues. Among these, *Gata3* emerged as a representative epigenetically repressed gene in normal lung tissue: HDM exposure led to consistent promoter hypermethylation and corresponding downregulation across biological replicates (hyper-down class; **Figure 6C**), consistent with suppression of Th2 signaling. This repression of *Gata3* by HDM is surprising, as HDM exposure is typically associated with promoting Th2 responses, for which GATA3 is a key transcriptional regulator. In tumors, *Gata3* expression was already low at baseline and showed minimal change with HDM treatment, indicating tumor-intrinsic repression.

Collectively, these data suggest that while HDM exposure can alter immune gene expression and epigenetic regulation in normal lung tissue, the TME is already epigenetically reprogrammed to suppress key immune pathways, limiting its responsiveness to further environmental cues.

### IL-17A, but not IL-1β, is required for HDM-driven lung tumor promotion

Given our transcriptomic and epigenetic data implicating IL-1 and IL-17 signaling pathways in HDM-induced immune remodeling, we next assessed their functional roles in tumor progression. LLC tumors were implanted into *Il1b*^−/−^ and *Il17a*^−/−^ mice using the same HDM exposure protocol as in WT mice **(Figure 7A)**. The pro-tumorigenic effect of HDM was preserved in *Il1b*^−/−^ mice **(Figures 7B–D)**, indicating that IL-1β is dispensable despite transcriptional enrichment of IL-1 signaling and inflammasome components in HDM-exposed normal lung tissue. In contrast, HDM-mediated tumor promotion was abolished in *Il17a*^−/−^ mice **(Figures 7E–G)**, highlighting the essential role of IL-17A. VEH-treated *Il17a*^−/−^ mice also exhibited reduced tumor multiplicity and tumor area compared to WT controls **(Figure 1)**, despite receiving equal numbers of viable tumor cells at implantation, indicating that IL-17A contributes to both tumor seeding and growth in the LLC model, consistent with previous reports^21–24^.

**Figure 7.**
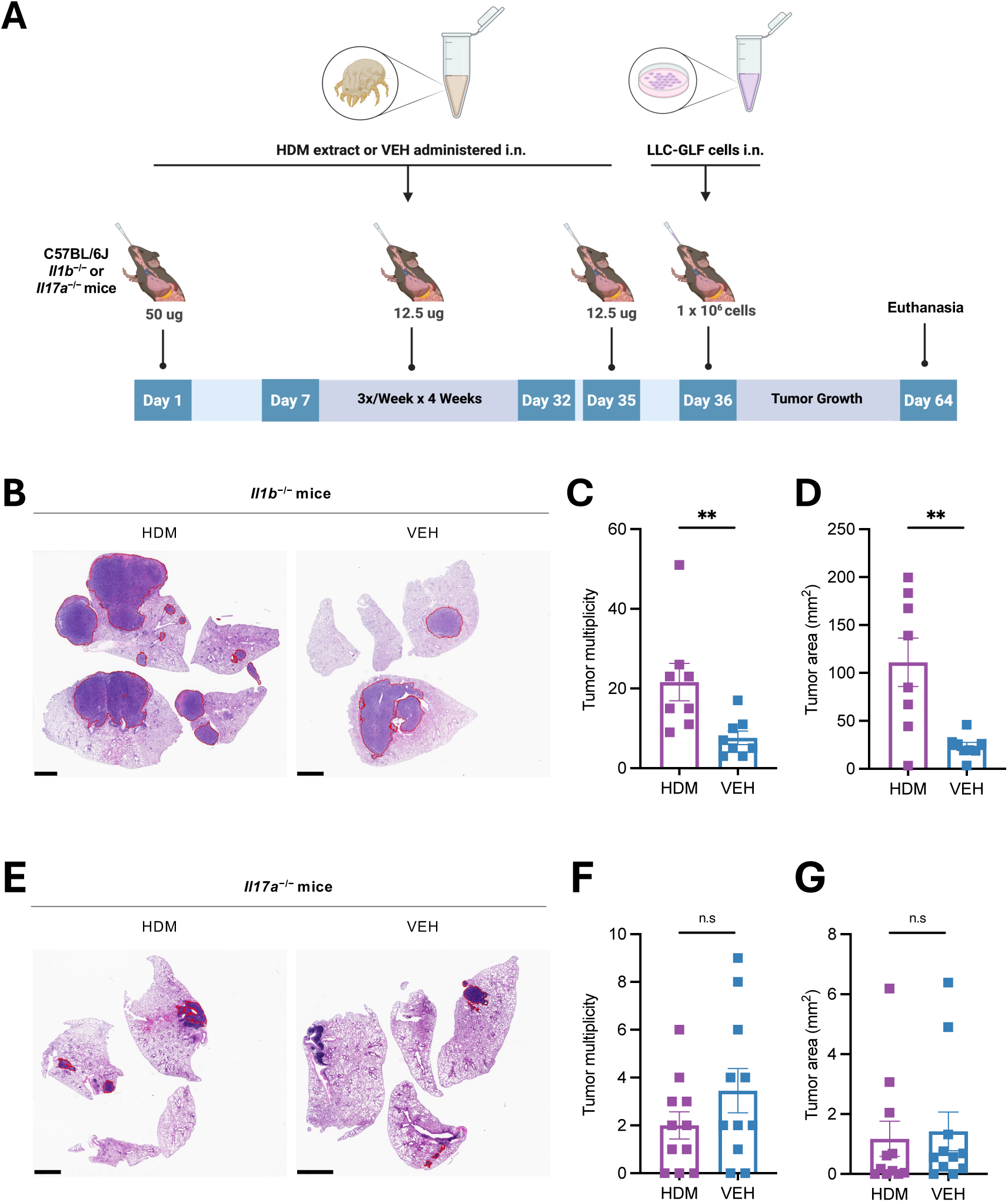
HDM-Driven Lung Tumor Promotion Requires IL-17A but Not IL-1β. **A)** Age- and sex-matched C57BL/6J *Il1b*^−/−^ (n = 8 mice/group) or *Il17a*^−/−^ mice (n = 11 mice/group) were first sensitized by i.n. instillations under light isoflurane anesthesia with HDM (50 µg/50 µL per mouse) or vehicle control (VEH; 50 µL per mouse) on Day 0, followed by three i.n. challenges per week (M/W/F) for four weeks with HDM (12.5 µg/50 µL per mouse) or VEH (50 µL/mouse). LLC-GLF cells (1 × 10^6^) were administrated i.n. on Day 36, 24h after the last HDM or VEH i.n. challenge. Mice were euthanized on Day 64, and the lungs were harvested for analysis as outlined in the schematic overview of the study design. **B)** Representative photos of H&E-stained lung sections of *Il1b*^−/−^ treated with either HDM or VEH. The areas within the red borders were considered positive for lung tumors. Scale bars, 2 mm. **C)** Tumor multiplicity calculated on H&E-stained sections as shown in B. **D)** Tumor area calculated on H&E- stained sections as shown in B. Data are presented as mean ± SEM. **E)** Representative photos of H&E-stained lung sections of *Il17a*^−/−^ treated with either HDM or VEH. The areas within the red borders were considered positive for lung tumors. Scale bars, 2 mm. **F)** Tumor multiplicity calculated on H&E-stained sections as shown in E. **G)** Tumor area calculated on H&E-stained sections as shown in E. Data are presented as mean ± SEM. Statistical significance was assessed by Mann-Whitney test; n.s: non-significant, and ** *p* <0.01.

These findings align with our RNA-seq and module score analyses: HDM exposure robustly activated IL-17 signaling in normal lung tissue but suppressed T cell responses in tumors. IL-17 likely shapes a lung microenvironment enriched in suppressive myeloid cells and depleted of cytotoxic T and NK cells, thereby facilitating immune evasion and tumor progression^25,26^.

Collectively, these data demonstrate that chronic HDM exposure promotes lung cancer progression via IL-17-associated pathways, positioning IL-17A as a central mediator of HDM-induced tumor promotion.

## Discussion

How environmental exposures, such as the aeroallergen HDM, influence lung cancer development remains poorly understood. Chronic lung inflammation induced by aeroallergens may remodel the tumor immune microenvironment and accelerate lung tumor initiation and progression, with implications for immunotherapy and precision oncology. To address this gap, we applied a multi-omics approach to investigate the effects of HDM exposure on lung tumor development in mice.

Previous studies have shown that HDM stimulates ROS production, leading to genotoxic stress and DNA damage in airway epithelial cells, including the BEAS-2B cell line^3,4^. HDM can also downregulate OGG1, a critical enzyme that excises oxidized guanine bases (8- oxoG) from DNA, thereby impairing the repair of oxidative lesions^27,28^. Based on these findings, we hypothesized that HDM may act as a mutagen and induce somatic mutations that accelerate lung cancer development. To test this hypothesis, we first evaluated the mutagenic potential of HDM *in vitro* using human BEAS-2B cells, which harbor wild-type p53 and proficient DNA repair^29^, and mouse LLC cells, which carry mutant p53 and exhibit genomic instability, including defects in mismatch repair and p53-dependent responses^30^. Under the tested conditions (a single 24-hour exposure), HDM did not induce detectable mutagenesis in either cell type, regardless of p53 status or DNA repair capacity. Similarly, chronic HDM exposure *in vivo* did not induce new mutations or significantly increase the mutational burden in tumors or normal adjacent tissues, despite a trend toward higher mutational burden in HDM-treated samples.

Although HDM was not directly mutagenic *in vitro*, it may indirectly promote mutagenesis *in vivo* through inflammation and oxidative stress, creating a pro-mutagenic microenvironment. To explore this possibility, we examined the effects of chronic HDM exposure in an orthotopic lung tumor mouse model. Consistent with our prior studies in other lung cancer models^31^, chronic HDM exposure significantly accelerated lung tumor formation and growth. However, even after prolonged exposure, HDM did not induce new mutations or significantly increase the overall mutational burden in either tumors or adjacent normal lung tissue, although a trend toward higher mutation counts in HDM-treated samples was observed. These data suggest that HDM’s pro-tumorigenic effects arise primarily through non-mutagenic mechanisms. Indeed, our multi-omics analyses point toward a mechanism involving immune and microenvironmental reprogramming. HDM exposure elicited divergent effects in normal lung tissue and lung tumors, with a bias towards affecting the lung microenvironment more than the tumor. In normal adjacent lung tissue, HDM triggered a robust pro-inflammatory response, affecting both innate (e.g., neutrophil degranulation, FCGR signaling) and adaptive immune pathways (e.g., B cell receptor signaling, interleukin-related genes), suggesting that allergen exposure primed the lung microenvironment for heightened immune activity. In contrast, within tumors, HDM suppressed pathways critical for antitumor immunity, including chemokine signaling, T cell differentiation, and antigen presentation, suggesting that tumors actively reprogram HDM-induced inflammation into an immunosuppressive state that facilitates immune escape. In addition to immune pathway changes, HDM exposure altered oxidative phosphorylation and respiratory electron transport in both tumors and normal lung tissue, indicating altered mitochondrial respiration. These metabolic shifts are consistent with a Warburg-like reprogramming that can support tumor growth and immune evasion, possibly by altering nutrient availability or redox signaling in the TME.

These findings are supported by immune deconvolution analysis, which indicated a shift toward a myeloid-dominant TME with monocyte enrichment and reduced NK cell and CD8^+^ T cell infiltration in tumors from HDM-exposed mice. This immune landscape is characteristic of an immunosuppressive TME and is associated with poor antitumor immune surveillance. Our integrative analysis of DNA methylation and transcriptomics further supports this immunosuppressive model. Among the 20 genes under apparent “epigenetic control” in normal tissue, several were immune-related, including *Gata3*, a key Th2 transcription factor, which was hypermethylated and downregulated following HDM exposure. While HDM is known to induce Th2 responses, chronic exposure may trigger complex immune regulation, including epigenetic repression of *Gata3* as a potential feedback mechanism to limit excessive inflammation. This repression could also reflect tissue-specific effects in epithelial cells, distinct from effects on immune cells, and aligns with the role of *Gata3* in tumors, where it is frequently suppressed to facilitate immune evasion^32^. Indeed, *Gata3* expression was already low in tumors at baseline and was further downregulated with HDM exposure, indicating both tumor-intrinsic repression and HDM-induced suppression. In line with this, promoter methylation patterns in tumors were comparatively static, suggesting that the malignant epigenome is preconfigured to silence key immune functions. This apparent lack of methylation plasticity in tumors may reflect pre-existing promoter hypermethylation or chromatin remodeling events that establish and maintain immune-suppressive transcriptional programs.

Since IL-1 and IL-17 signaling pathways were activated as part of HDM-induced immune remodeling and are generally considered pro-tumorigenic in lung cancer^25,26,33^ we examined the impact of *Il1b* and *Il17a* gene deletion in our model. In prior work, we found that IL-1β neutralization suppressed the pro-tumorigenic effects of HDM in a urethane-induced model and in *Kras^G12D^*-genetically engineered mice^31^. However, in the present study, deletion of *Il1b* did not alter the tumor-promoting effects of HDM in the LLC model. This suggests that IL-1β signaling contributes to early disease stages (e.g., hyperplasia and adenoma in urethane-treated and *Kras^G12D^* models), but is dispensable in advanced disease (adenocarcinoma in the more aggressive LLC model). This interpretation is consistent with recent clinical trials of Canakinumab (an IL-1β-neutralizing antibody) that failed to show efficacy in patients with locally advanced or metastatic non-small cell lung cancer^34^. A possible explanation is that IL-1β signaling, while pro-tumorigenic in some contexts, may also be required to sustain antitumor T cell responses^33,35^. In contrast, blockade of the IL-17 pathway has demonstrated consistent antitumor effects across numerous preclinical models^21–26,33^. In line with these studies, we found that lung tumor development was reduced in the absence of IL-17A, and HDM-driven tumor promotion was completely abrogated. These results identify IL-17A as a key mediator of tumor progression, both at baseline and under HDM exposure, thereby reinforcing the rationale for targeting the IL-17 axis in lung cancer.

In conclusion, our findings reveal that chronic HDM exposure fosters a pro-tumorigenic lung microenvironment primarily through immune modulation, metabolic reprogramming, and selective epigenetic remodeling, rather than direct mutagenesis. We propose a model in which HDM primes the lung with an inflammatory microenvironment, while tumors reshape these signals into an immunosuppressive, myeloid-skewed phenotype that supports immune escape. These insights suggest that chronic allergic airway inflammation may accelerate lung cancer development and highlight the therapeutic potential of targeting IL-17A or reconditioning the immune microenvironment in environmentally influenced cancers.

### Limitations of the study

Although our study did not identify a mutagenic effect of HDM, the limited sample size may have reduced the statistical power and contributed to the absence of significant differences between HDM- and VEH-treated conditions for parameters such as the mutational burden. A larger sample size may have yielded different results. Additional factors, such as the single 24-hour *in vitro* exposure and the timing of tumor cell inoculation relative to the HDM exposure period *in vivo*, may also have limited the detection of mutagenic effects. Future studies using models with impaired DNA repair, or alternative HDM exposure protocols may reveal conditions under which HDM exerts mutagenic activity. Our study also focused on a single aeroallergen. Other environmental exposures, such as fungi (e.g., *Aspergillus*, which can induce IL-17-mediated airway inflammation^36^ or pollens (e.g., Ragweed, which can cause DNA damage^37^, may promote mutagenesis and tumorigenesis. Future work should investigate these additional aeroallergens. Finally, the clinical relevance of our findings remains to be established. Whether long-term HDM exposure represents a risk factor for lung cancer in humans warrants investigation, particularly in genetically predisposed individuals (e.g., those harboring oncogenic mutations) or in individuals co-exposed to other lung carcinogens.

## Experimental model and study participant details

### Mice

C57BL/6J WT mice were purchased from the Jackson Laboratory (JAX, Strains No. 000664). Initial breeding pairs *Il1b^−/−^* mice (JAX, Strain# 034447) backcrossed to C57BL/6J for over 10 generations were kindly provided by Dr. Wai Wilson Cheung, Dr. Robert Mak, and Dr. Hal Hoffman (UC San Diego). Initial breeding pairs of *Il17a^−/−^* mice on the C57BL/6J background were kindly provided by Dr. Rachel Caspi (NIH). All mice were bred in our vivarium under specific pathogen-free (SPF) conditions for more than 6 months and knockout strains were genotyped before being used in experiments. Mice were kept on a 12-hour light and 12-hour dark cycle with ad libitum access to standard chow and water. Both male and female mice were used in this study. Mice were randomly assigned to experimental groups based on genotype, treatment, and matched for age and sex to minimize bias. Mice were monitored periodically, and any animals showing signs of significant distress, such as a body weight (BW) loss >20% of initial BW, hunching posture, inability to eat or drink, or moribundity, were removed from the study and humanely euthanized. Mice that did not survive until the study endpoint were excluded, and this attrition was accounted for in the analysis. All the animal studies were approved by the Institutional Animal Care and Use Committee (IACUC) at the University of California San Diego (Protocol Number: S02240) and were conducted in accordance with the approved protocols, adhering to the Animal Research: Reporting of In Vivo Experiments (ARRIVE) guidelines.

### Cells and cell culture

Human bronchial epithelial cells (BEAS-2B) were purchased in 2018 from ATCC (Cat. No. CRL-9609). Lewis lung carcinoma (LLC) cells^6^ and Luciferase-expressing LLC (LLC- GLF) cells^38^ were obtained from Dr. Jack Bui and Dr. Tomoko Hayashi (UC San Diego), respectively. LLC-GLF cells were authenticated via short tandem repeat (STR) profiling by IDEXX BioResearch. The STR profile matched identically to the genetic profile established for the LLC cell line (ATCC, Cat. No. CRL-1642). BEAS-2B and LLC cell lines were cultured under standard conditions and used for experiments within five passages after thawing. Briefly, BEAS-2B cell stocks were maintained in complete Bronchial Epithelial Growth Medium (BEGM; Lonza, Cat. No. CC-3170). For experimental procedures, BEAS-2B and LLC cells were cultured in Roswell Park Memorial Institute (RPMI) 1640 medium (Gibco, Cat. No. 11875-093) and Dulbecco’s Modified Eagle Medium (DMEM; Gibco, Cat. No. 11995-065), respectively. Both media were supplemented with 10% fetal bovine serum (FBS; Gibco, Cat. No. 26140079) and 1% penicillin-streptomycin (Gibco, Cat. No. 15140122). The cells were maintained in a humidified incubator at 37°C with 5% CO₂. Culture media were refreshed every 2-3 days, and cells were passaged at ∼80% confluence using 0.25% Trypsin-EDTA (Gibco, Cat. No. 25200072). Both cell lines were tested for bacterial and fungal contamination using a cell culture contamination detection kit (Thermo Fisher Scientific, Cat. No. C7028) prior to their use in experiments.

## Method details

### HDM extract solubilization

Lyophilized extracts from whole bodies of HDM *Dermatophagoides pteronyssinus* species (Greer laboratories, Cat. No. XPB82D3A25) were reconstituted in sterile PBS (Thermo Fisher Scientific, Cat. No. 14190144) at a concentration of 10 mg/mL (based on their total protein content), aliquoted, and stored at -80 °C until used.

### *In vitro* HDM exposure model

BEAS-2B cells and LLC cells were used to evaluate the potential mutagenic effects of HDM exposure. HDM concentrations were determined based on the IC_50_ values for each cell line, corresponding to concentrations inducing approximately 50% cell death. BEAS- 2B cells were seeded at 5.0 × 10⁴ cells per well, and LLC cells at 2.5 × 10⁴ cells per well, in separate 24-well plates and left adhere overnight. Cells were treated with either 200 µg/mL HDM, 400 µg/mL HDM, or the same volume (50 µL) of PBS as negative control. After 24 hours of exposure, cells were washed in PBS and cultured for another 24 hours in their respective growth media. Cells were then expanded first into T25 flasks, and subsequently into T75 flasks upon reaching confluency. Cells were harvested and aliquots of 1.0 × 10⁶ cells were used for DNA and RNA extraction.

### HDM IC₅₀ determination

The half-maximal inhibitory concentration (IC₅₀) of HDM on BEAS-2B and LLC cells was determined using the CellTiter-Glo^®^ Luminescent Cell Viability Assay (Promega, Cat. No. G7570). BEAS-2B cells were seeded at 10,000 cells per well and LLC cells at 20,000 cells per well in white 96-well plates (Corning, Cat. No 3610) and allowed to adhere overnight. Cells were then treated with a range of HDM concentrations (0-1000 µg/mL) in triplicate for 24 hours. Following exposure, CellTiter-Glo^®^ reagent was added 1:1 to each well, incubated for 10 minutes at room temperature, and luminescence was measured using a microplate reader (BioTek Synergy or equivalent). Raw luminescence values were normalized to PBS-treated controls, and dose-response curves were generated using nonlinear regression in GraphPad Prism v10.6.0. IC_50_ values were defined as the HDM concentration causing a 50% reduction in luminescence relative to PBS control.

### *In Vivo* HDM exposure model

WT C57BL/6J mice were sensitized by intranasal (i.n.) instillation with HDM (50 µg/50 µL per mouse) or VEH control (PBS; 50 µL per mouse) under light isoflurane anesthesia on Day 0, followed by three i.n. challenges per week for four weeks with HDM (12.5 µg/50 µL per mouse) or PBS (50 µL per mouse). On Day 32, mice were orthotopically implanted with 1 × 10⁶ LLC-GLF cells via i.n. administration, delivered in two separate doses of 0.5 × 10⁶ cells in 50 μl PBS, one in the morning and one in the afternoon, to minimize variability in delivery efficiency and reduce the risk of administration error. Tumor growth was monitored weekly by bioluminescence imaging (BLI) using an *in vivo* imaging system (IVIS), as previously described^7^. Briefly, hair was removed from the throat to the lower abdomen using a hair removal cream, and D-luciferin (GOLDBIO, Cat. No. LUCK-100) was injected intraperitoneally (i.p.) at 150 mg/kg in sterile PBS approximately 10 to 15 minutes before each BLI session. On Day 64, 24h after the last BLI, mice were euthanized by CO2 asphyxiation, and the lungs were harvested for subsequent analyses.

### Histological analysis

Four lung lobes were fixed in 10% buffered formalin for 24h and brought in histological grade 70% ethanol to the UC San Diego Moores Cancer Center Tissue Technology Shared Resource for paraffin embedding and sectioning. The fixed lungs were cut into 4- 6-µm sections, placed on glass slides, and stained with hematoxylin (Thermo Fisher Scientific, Cat. No. 7221) and eosin (Thermo Fisher Scientific, Cat. No. 7111) (H&E) on a Gemini AS slide stainer (Thermo Fisher Scientific) using standard staining procedures. H&E slides were scanned on an Aperio AT2 slide scanner (Leica Biosystems), digitized, and whole-slide images were used for tumor assessments. Tumor measurements were performed in a blinded fashion using QuPath software version 0.5.1^8^. The tumor multiplicity (i.e., the sum of lung lesions per mouse) was calculated on one H&E-stained section of 4 lung lobes for each mouse. The tumor area for each mouse was determined by summing the surface areas of all lung tumors and was expressed in mm^2^ as previously described^31^.

### DNA extraction from cell lines

Genomic DNA was extracted from cultured cells using the DNeasy Blood & Tissue Kit (Qiagen, Cat. No. 69504), following the Animal Blood or Cells Spin-Column protocol according to manufacturer’s instructions.

### DNA and RNA extraction from tissues

Macroscopic LLC tumors and normal adjacent lung tissues were manually dissected using a binocular microscope and a scalpel. The tissues were snap-frozen in liquid nitrogen and stored at -80 °C until use. The samples were then sectioned into ∼1 mg aliquots for nucleic acid extraction. Genomic DNA and total RNA were co-extracted using the AllPrep DNA/RNA Mini Kit (Qiagen, Cat. No. 80204). Tissue was homogenized with PowerTube beads (Cat. No. 13117-50) using the Omni Bead Ruptor 12 (Cat. No. 19- 050A). DNA concentration was quantified using the Qubit™ 1X dsDNA HS Assay Kit (Cat. No. Q33231) on the Qubit 4 Fluorometer (Cat. No. Q33226). All DNA and RNA samples had a DNA Integrity Number (DIN) and RNA Integrity Number (RIN), respectively, greater than 6.

### DNA sequencing

Whole-genome Universal Duplex sequencing (UDseq) at 90X was performed for normal lung tissues and cell lines. Two hundred ng of input DNA was enzymatically fragmented using dsDNA Fragmentase (New England Biolabs, Cat. No. M0348S) for 20 minutes to achieve ∼300 bp fragments. Libraries were prepared using the cfDNA & FFPE DNA Library Prep v2 MC Kit (Roche, Cat. No. 10010206). Adapter-ligated molecules were quantified via qPCR and indexed using PCR amplification with i7 and i5 primers. Fragment size distributions were assessed using the Agilent 4150 TapeStation System (Cat. No. G2992AA) with the HS D5000 ScreenTape (Cat. No. NC1874887) and HS D5000 Reagents (Cat. No. 5067-5589). Bulk whole genome sequencing (WGS) at 90X was performed for tumor tissues and at 30X for the spleen samples. Two hundred ng of DNA was fragmented with dsDNA Fragmentase (Cat. No. M0348S) for 20 minutes. Libraries were prepared using the NEBNext Ultra II End Repair/dA-Tailing Module (Cat. No. E7546L), the NEBNext Ultra II Ligation Module (Cat. No. E7595L) and indexed using i7 and i5 primers. DNA libraries were submitted to Novogene (Sacramento, CA) and sequencing on Illumina NovaSeq X-Plus Sequencing Platform.

### DNA sequencing data analysis

To identify high-confidence somatic mutations, we utilized a duplex sequencing approach, which allows for the accurate detection of ultra-rare mutations by independently tagging and sequencing both strands of each DNA molecule. Raw sequencing data (FASTQ files) were first aligned to the appropriate reference genomes (GRCm39 for mouse samples and GRCh38 for human samples) using BWA-MEM (v0.7.17). Duplicate reads were marked and removed using Picard tools, and alignment files were sorted and indexed with SAMtools. For variant calling, we applied an ensemble approach using four independent somatic variant callers: Mutect2, VarScan2, Strelka2, and MuSe. Variants identified by at least two of the four callers were retained as candidate somatic mutations. To further increase specificity, we applied the following post-calling filters: dbSNP annotation (via Variant Effect Predictor) was used to remove known germline single nucleotide polymorphisms (SNPs). Shared mutations across multiple samples and clustered mutations (defined as > 3 mutations within a 10 bp window) were excluded to eliminate potential germline contamination or regions under positive selection.

### RNA sequencing

Total RNA samples were submitted to Novogene (Sacramento, CA) for library preparation and paired-end sequencing. Post-library quality control was performed using Agilent High Sensitivity D1000 ScreenTape. Sequencing was conducted on the Illumina NovaSeq X- Plus Sequencing Platform. All samples achieved high-quality metrics, including Q30 scores >90% from Illumina sequencing, and sequencing saturation >80% and mean mapping rate of 98% to the Mus musculus reference genome (GRCm39/mm39).

### RNA sequencing data analysis

Twenty million 150-bp paired-end reads were generated on the Illumina (insert here), and primary sequence data was then processed using the Illumina DRAGEN BCL Convert 07.021.645.4.0.3 pipeline. Raw sequencing data were trimmed to remove low-quality reads using Trim Galore. Cleaned sequence reads were then aligned against the *Mus musculus* genome (GRCm39/mm39) using the STAR aligner (v2.3.5a). Gene-level read counts were generated from the aligned data. Only primary alignments with mapping quality ≥10 were retained. Gene-level counts were obtained with featureCounts from exonic regions, summarized by gene_id and treating paired-end reads as fragments. Differential expression was performed with DESeq2. Genes with very low counts were filtered (e.g., row sum <10). Size factors were estimated by the median-of-ratios method, dispersions were modeled per gene, and Wald statistics were used for inference with the design ∼ Tissue + Treatment + Tissue:Treatment (or the exact design you used). *p*-values were adjusted by Benjamini-Hochberg FDR; genes with FDR < 0.05 were considered significant. Pathway enrichment analysis was performed using ConsensusPathDB (CPDB)^15^, integrating annotations from the KEGG, Reactome, and MouseCyc pathway databases. Pathways identified using the default database annotations, without manual curation or filtering, are presented in the Supplemental Information. Curated pathways, emphasizing disease- and tissue-specific processes, are shown in the main figure. Pathways were considered significantly enriched if they contained at least 3-5 genes with a *p*-value ≤ 0.05. To account for multiple hypothesis testing, pathways with a *q*-value > 0.05 were excluded from further analysis.

### DNA methylation profiling

One µg of genomic DNA was used for DNA methylation profiling with the Infinium Mouse Methylation BeadChip Kit v1.0 (Illumina, Cat. No. 20041558). BeadChip processing, bisulfite conversion, array hybridization, and signal scanning using the Illumina iScan System were performed according to the manufacturer’s instructions at the UC San Diego IGM Genomics Center.

### DNA methylation data analysis

Genome-wide DNA methylation was analyzed from Illumina Infinium raw IDAT files. Two cohorts were analyzed separately (Tumor and Normal tissue), each with three biological replicates per condition (HDM, n = 3; VEH, n = 3). IDAT files were processed in R using RnBeads^39^, using default settings unless otherwise specified. The following options were explicitly applied for both cohorts: probes on sex chromosomes were removed (filtering.sex.chromosomes.removal = TRUE), sample identities were mapped via the Sample_ID field in the annotation (identifiers.column = "Sample_ID"), data were normalized using the wm.dasen method (normalization.method = "wm.dasen") together with methylumi background correction (normalization.background.method = "methylumi.noob"). Analyses were performed against the mm10 genome assembly (assembly = "mm10", via RnBeads.mm10). Group comparisons were defined by the annotation column Treatment (e.g., HDM vs VEH; differential.comparison.columns = "Treatment"), and site-level differential reports were generated (differential.report.sites = TRUE). Results were exported as CSV files (export.to.csv = TRUE) for the following feature classes: tiling regions, gene bodies, promoters, and CpG islands. Differentially methylated sites were summarized as counts of hyper- and hypomethylated loci by tissue (HDM vs VEH) across tissue types and all genomic regions (genes, promoters, CpG islands, tiling), using a significance threshold of |fold change| ≥ 0.20 and *p* < 0.05. Per-probe beta values were obtained from RnBeads exports. Promoter annotations were used to map probe/region identifiers to gene symbols. Methylation and RNA-Seq Integration To evaluate the relationship between promoter methylation and gene expression in HDM treated samples in two tissue types, we extracted common genes present in both datasets and constructed two matrices: one containing methylation values and the other containing expression values. Each matrix consisted of three HDM samples aligned by gene identifiers. Genes with missing values in either dataset were excluded to ensure complete cases across all samples. For each gene, we calculated correlations between methylation and expression levels using two-sided statistical tests. Note, the gene expression matrix consisted of trimmed mean of M values (TMM) normalized RNA-seq values and the beta value matrix consisted of RnBeads normalized values. Linear associations were assessed with the Pearson correlation coefficient. Genes were prioritized based on strong anti-correlation between methylation and expression, defined as Pearson correlation coefficient r ≤ -0.6 with a *p*-value < 0.1. Given the limited sample size (n = 3), FDR correction did not yield significant results, and correlations were interpreted with caution. All analyses were performed in R (base functions cor.test and p.adjust) to ensure reproducibility. Emphasis was placed on effect size (magnitude of correlation) combined with FDR thresholds, and robust candidates were highlighted in the figures. For both normal and tumor tissues, promoter methylation data were integrated with RNA-seq differential expression results by mapping gene symbols to Ensembl identifiers and retaining genes with valid annotations and significant promoter methylation (*p* < 0.05). For each gene, the log2 fold change from RNA-seq was paired with the mean.quot.log2 methylation difference, and genes were classified into five categories based on effect size thresholds: Hyper-Up (hypermethylated, upregulated), Hyper-Down (hypermethylated, downregulated), Hypo-Up (hypomethylated, upregulated), Hypo-Down (hypomethylated, downregulated), and Low effect size. Classification cutoffs were defined as |log2 fold change | ≥ 1 for RNA-seq and |log2 fold change | ≥ 0.1 for methylation. Genes were visualized using scatter plots in R (ggplot2), with methylation changes plotted on the x-axis and expression changes on the y-axis, color-coded by category and annotated with dashed lines to indicate cutoff thresholds; normal and tumor profiles were then displayed side-by-side for direct comparison.

### Immune deconvolution analysis

Bulk RNA-seq expression matrices (gene symbols × samples; unit: TMM-normalized log2 CPM counts) were input to mMCP-counter^14^. Differences in estimated cell fractions between groups (e.g., HDM vs VEH; Normal vs Tumor) were tested using two-sided Wilcoxon rank-sum tests and Benjamini-Hochberg FDR across cell types; effect sizes (median differences) are reported alongside adjusted *q*-values.

### Statistical analysis

Sample sizes for *in vivo* studies were determined based on preliminary data, observed variability in the lung cancer model, and power analysis. All animal studies were adequately powered to achieve statistically significant results while minimizing the number of animals used. The number of mice per group and the replication level for each *in vitro* and *in vivo* experiment are mentioned in the figure legends. Graphical representations were generated using R v4.5.1 or GraphPad Prism v10.6.0. Data are presented as mean ± SEM or median and interquartile range (IQR), as indicated. The statistical comparisons between two groups were performed using either a Mann-Whitney test or a two-sided Welch’s *t*-test, as specified in the figure legends. The statistical tests used for RNA-seq, immune deconvolution, DNA methylation, and DNA-seq analyses are detailed in the corresponding Methods sections. A *p*-value < 0.05 was considered statistically significant. All statistical analyses were conducted in R v4.5.1 or GraphPad Prism v10.6.0.

## Graphical illustrations

Graphical illustrations were created with BioRender.

## Resource Availability Lead contact

Requests for further information and resources and reagents should be directed to and will be fulfilled by the lead contact, Dr. Samuel Bertin.

## Materials availability

This study did not generate new unique reagents.

## Data and Code Availability

The raw FASTQ files and processed datasets generated in this study have been deposited in the GEO and SRA databases. Permanent accession numbers and links will be provided upon final acceptance and prior to publication. All analysis code will be made available on GitHub (https://github.com/shamsalazzam/code_Multi-omics-profiling), and all data and code will be publicly available without restriction.

## Acknowledgments

This study was supported by the US National Institutes of Health (NIH) grant U01CA276642 to E.R., N.J.G.W., and S.B.; a Targeted Grant from the Curebound Foundation (23TG03) to E.R. and S.B.; and by NIH grants R01ES032547, R01ES036931, R01CA269919, R01CA296974, P01CA281819, and U01CA290479 to L.B.A. Additional support was provided by L.B.A.’s Packard Fellowship for Science and Engineering and the UC San Diego Sanford Stem Cell Institute. Computational analyses were performed using the UC San Diego Triton Shared Computing Cluster (TSCC) at the San Diego Supercomputer Center (SDSC). We thank Dr. Valeria Estrada and Dr. Nissi Varki (UC San Diego) for guidance with histopathological evaluation of lung tumors; Dr. Jack Bui and Dr. Tomoko Hayashi (UC San Diego) for providing the LLC and LLC-GLF cell lines, respectively; the Biorepository and Tissue Technology Shared Resources (BTTSR) at the UC San Diego Moores Cancer Center, supported by the NCI CCSG grant P30CA23100, for tissue processing, embedding, H&E staining, and brightfield slide scanning; and Dr. Kristen Jepsen for guidance with the DNA methylation analysis, which was conducted at the UC San Diego IGM Genomics Center.

## Author contributions

Conceptualization, E.R., L.B.A., and S.B.; Methodology, S.A-A., I.S., S.S., M.Y-H., K.S., M.B., N.A-A., S.N., M.Z., J.D., and T.Y.; Validation, S.A-A., I.S., S.S., M.Y-H., and A-T.; Formal analysis, S.A-A., I.S., S.S., M.Y-H, A-T., N.A-A., T.Y., and N.J.G.W.; Investigations, S.A-A., I.S., S.S., and M.Y-H.; Resources, M.C., S.H.; Data curation: S.A-A., I.S., S.S., M.Y-H, and S.B.; Writing – original draft: S.A-A., and S.B.; Writing – review & editing: S.A-A., S.B., and L.B.A.; Visualization, S.A-A., I.S., S.S., and M.Y-H.; Supervision, E.R., L.B.A., and S.B.; Funding acquisition, L.B.A., S.B., N.J.G.W., and E.R.

## Declaration of Interests

L.B.A. is a co-founder, CSO, scientific advisory member, and consultant for io9 (now Acurion), has equity and receives income. The terms of this arrangement have been reviewed and approved by the University of California, San Diego in accordance with its conflict-of-interest policies. L.B.A. is a compensated member of the scientific advisory board of Inocras. L.B.A.’s spouse is an employee of Hologic, Inc. L.B.A. declares U.S. provisional applications filed with UC San Diego with serial numbers: 63/269,033; 63/289,601; 63/483,237; 63/412,835; 63/492,348; and 63/366,392 as well as a European patent application with application number EP25305077.7. L.B.A. and S.P.N. declare provisional patent application PCT/US2023/010679. L.B.A. is an inventor of a US Patent 10,776,718 for source identification by non-negative matrix factorization. All other authors declare that they have no competing interests.

---

**Figure S1.**
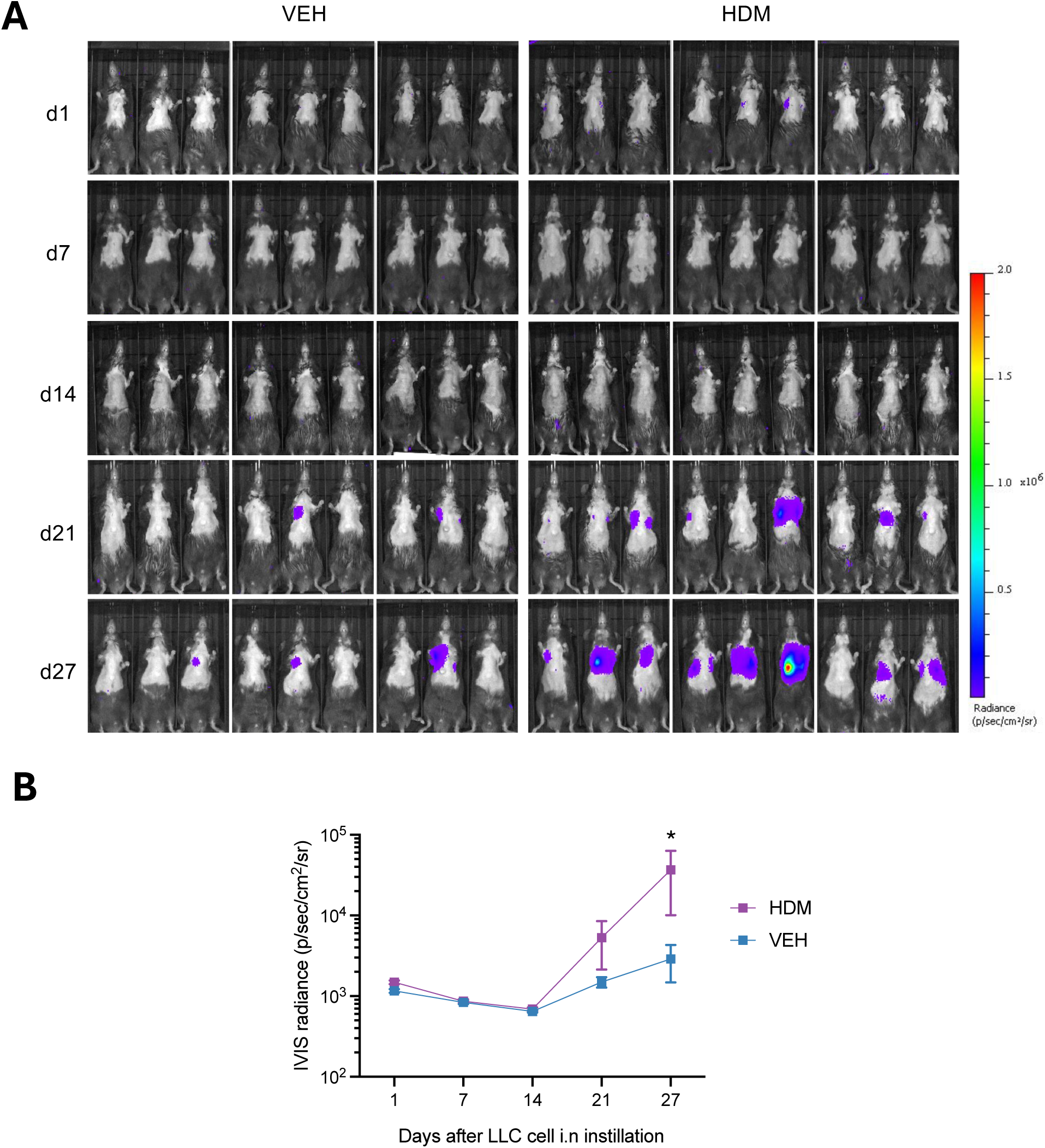
Kinetic of Tumor Growth in the LLC Orthotopic Lung Carcinoma Mouse Model. **A)** Representative IVIS images of tumor signals quantified and **B)** IVIS radiance in C57BL/6J WT mice (n = 9 mice/group) treated as shown in Figure 1A. Data are presented as mean ± SEM. Statistical significance was assessed by two-way ANOVA with post hoc Bonferroni’s test; * *p* <0.05.

**Figure S2.**
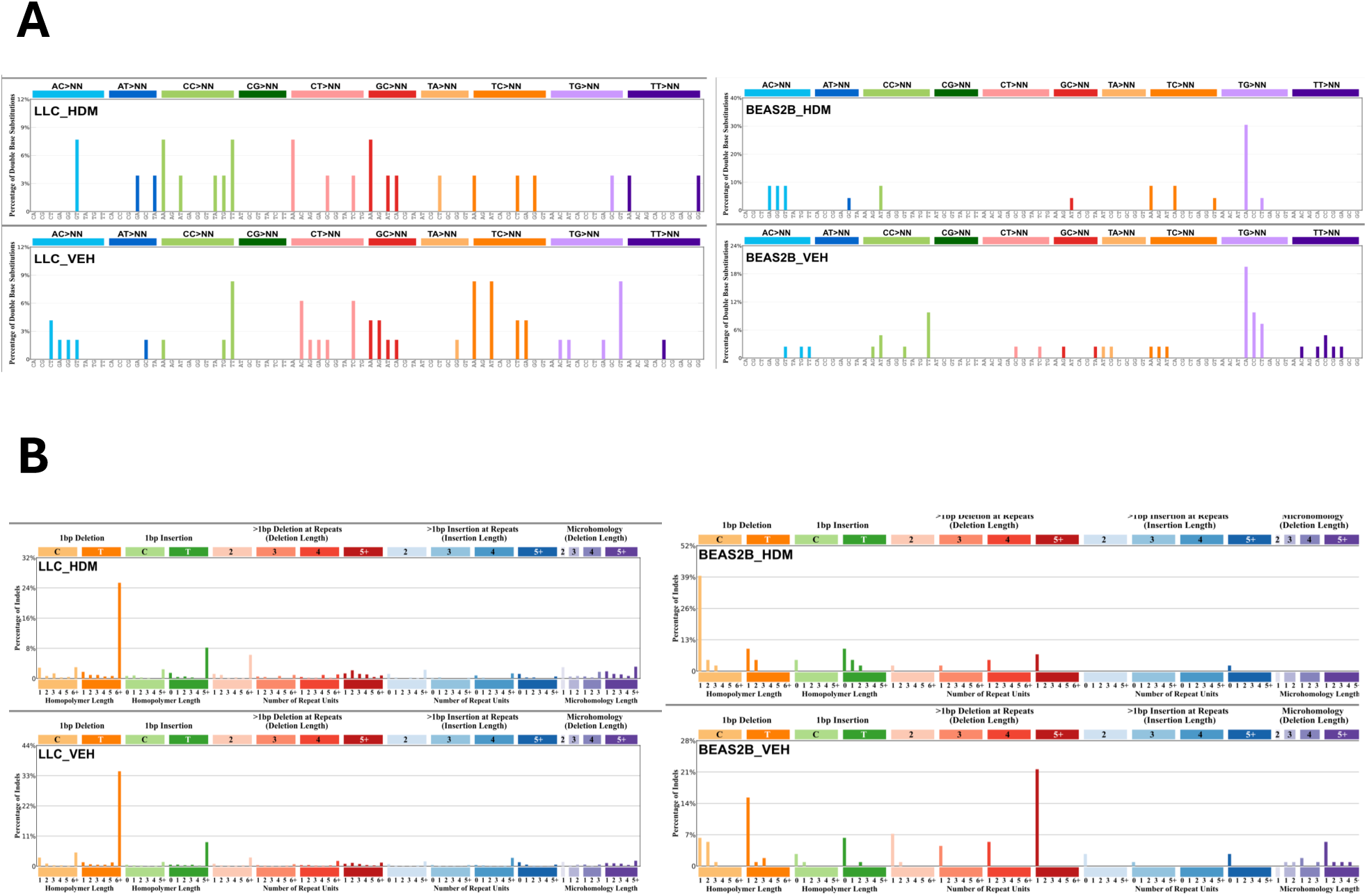
DBS78 and ID83 mutational profiles in HDM- and vehicle-treated LLC and BEAS- 2B cells. **A)** Double-base substitution signatures (DBS78) for HDM- and VEH-treated LLC and BEAS-2B cell lines. **B)** Insertion-deletion signatures (ID83) for the same samples and conditions as in A.

**Figure S3.**
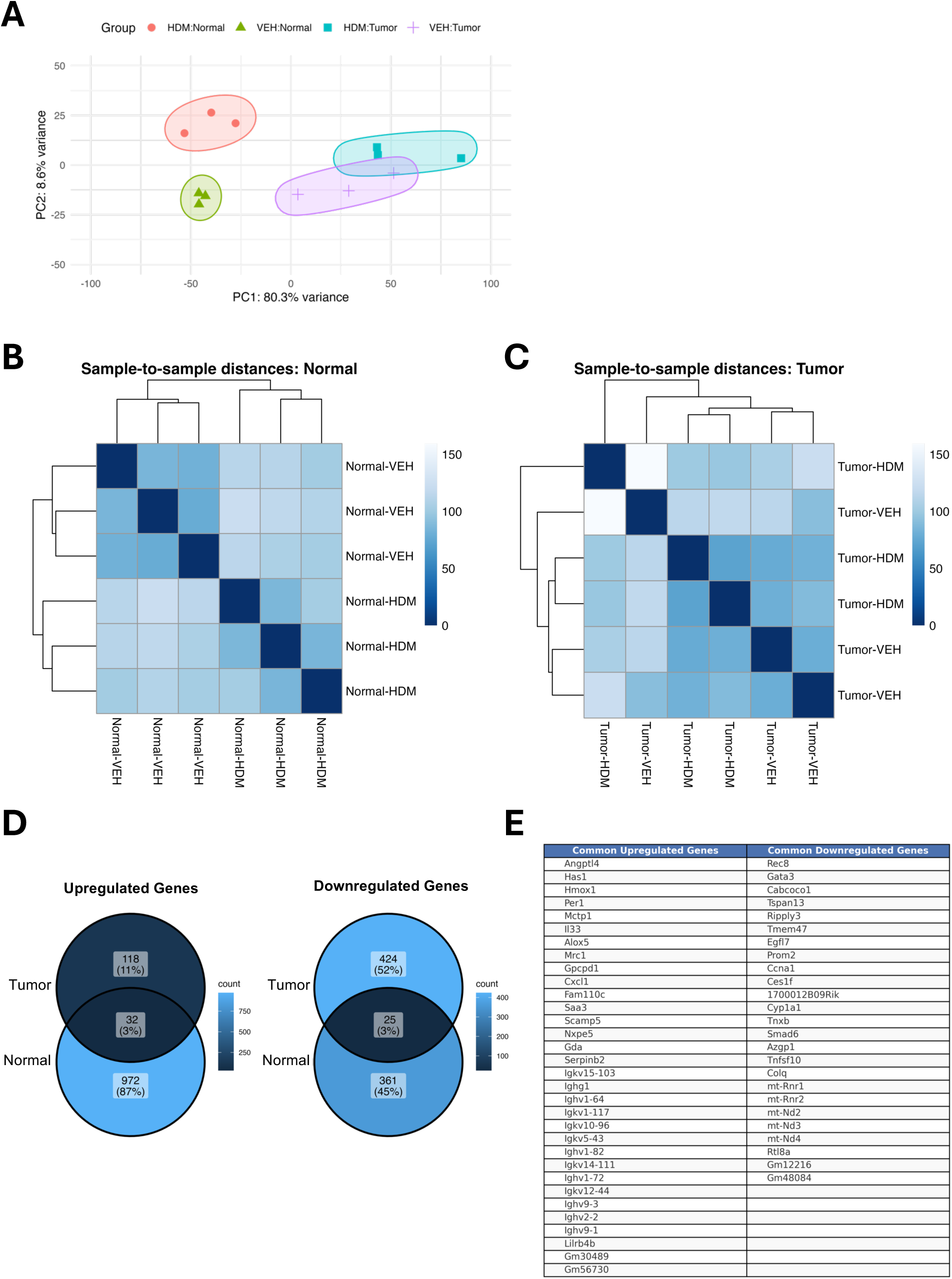
Sample Clustering in the RNA-seq Experiment. **A)** Principal component analysis (PCA) of samples. PCA was performed on log2(TPM+1) expression values (genes centered and scaled). Points are individual samples; color denotes exposure (HDM vs VEH) and shape denotes tissue (Normal vs Tumor). Ellipses show covariance-based group dispersion for visualization only (no hypothesis testing). **B)** Sample-to-sample distance heatmap of normal lung tissue, and **C)** of tumor samples. Pairwise Euclidean distances were computed from log2(TPM+1) expression and clustered by hierarchical clustering. The top dendrograms show clustering of samples; the left dendrograms show clustering of the distance profiles. Darker blue indicates smaller distances (greater similarity); lighter shades indicate larger distances (lower similarity). **D)** Venn diagrams showing overlap of upregulated (left) and downregulated (right) differentially expressed genes (DEGs) between normal and tumor tissues. Numbers indicate unique and shared gene counts, along with their relative percentages. **E)** Table listing DEGs shared between normal lung tissue and tumors, with upregulated genes on the left and downregulated genes on the right. Differential expression was assessed with a negative-binomial model (DESeq2, Wald test) with multiple testing controlled by Benjamini-Hochberg false discovery rate (FDR); genes with FDR < 0.05 were considered significant.

**Figure S4.**
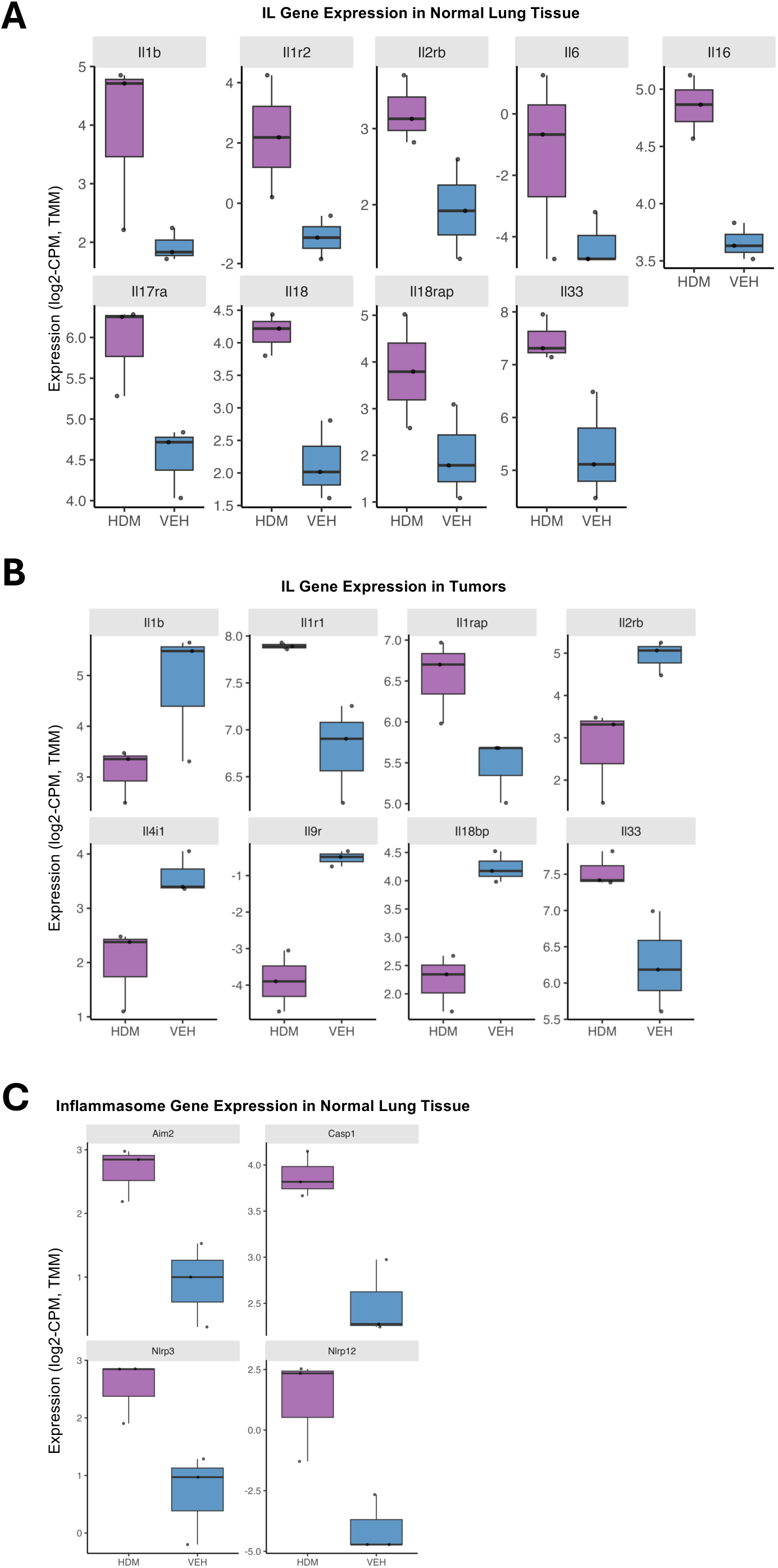
HDM Exposure Induces Distinct Immune Gene Signatures in Normal Lung Tissues and Tumors. **A)** Boxplots showing trimmed mean of M values (TMM)-normalized expression (log2-counts per million [CPM]) for selected interleukin (IL) cytokines and receptors in mice treated with HDM (purple) or VEH (blue) in normal lung tissue, and **B)** in tumors. Boxes indicate the median and interquartile range (IQR); whiskers extend to 1.5× the IQR; points are individual samples. **C)** Boxplots showing TMM-normalized expression (log2-CPM) for selected inflammasome-related genes in mice treated with HDM (purple) or VEH (blue) in normal lung tissue. Genes displayed were pre-selected as differentially expressed in the global analysis (DESeq2 Wald test with Benjamini-Hochberg FDR < 0.05).

**Figure S5.**
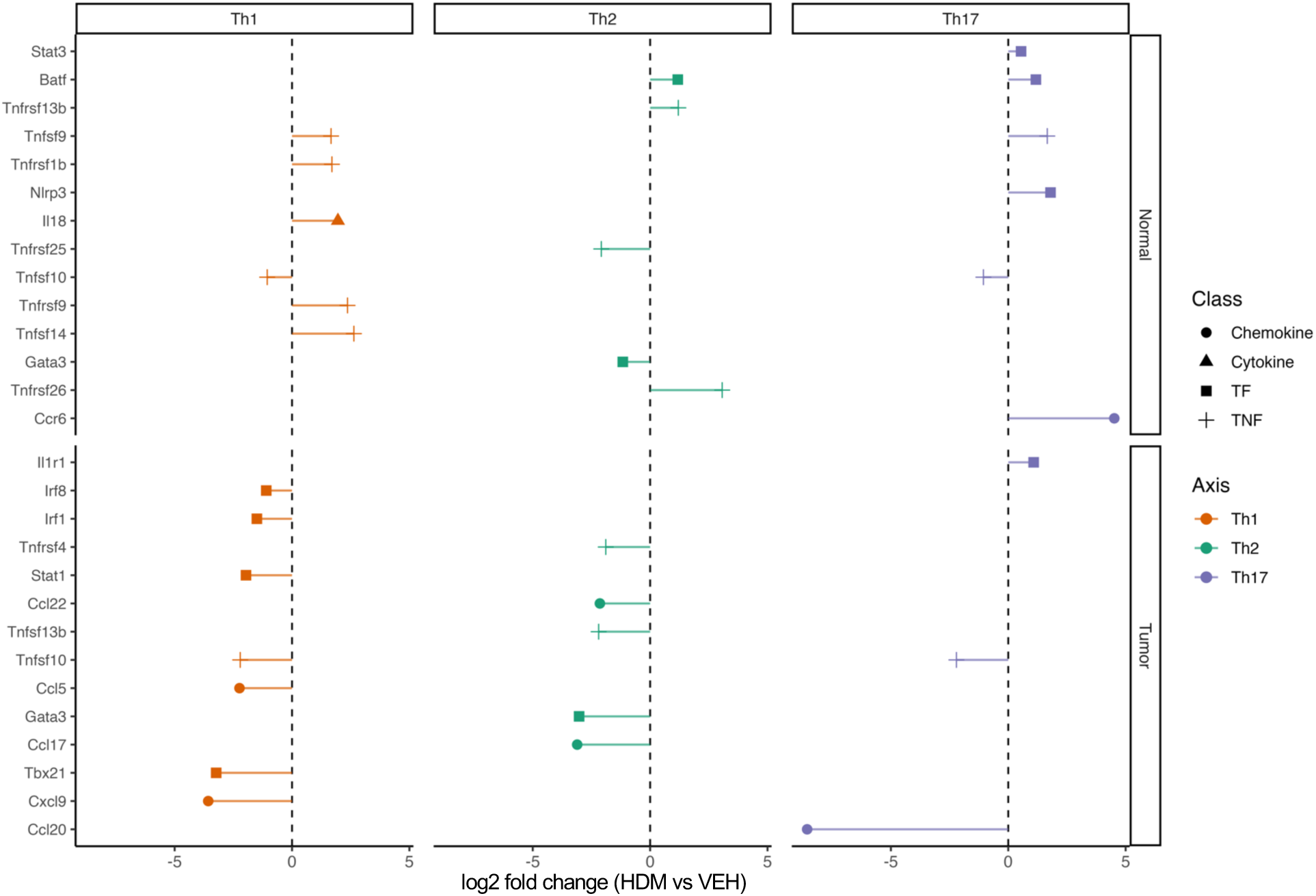
HDM Exposure Enhances T Cell Responses in Normal Lung Tissues but Suppressed Them in Tumors. Forest plots showing the log2 fold change (HDM vs VEH) for curated Th1, Th2, and Th17-associated genes in normal lung tissue (top) and tumors (bottom). Points represent the estimated log2 fold change for each gene. Point color encodes Th axis, and point shape denotes gene class: chemokine, cytokine, transcription factor (TF), or TNF family members. Differential expression was assessed per gene using a DESeq2 negative-binomial model with the Wald test (two-sided) on size-factor-normalized counts. *p* values were adjusted by Benjamini-Hochberg FDR in the global analysis; only genes with nominal *p* < 0.05 are shown.

**Figure S6.**
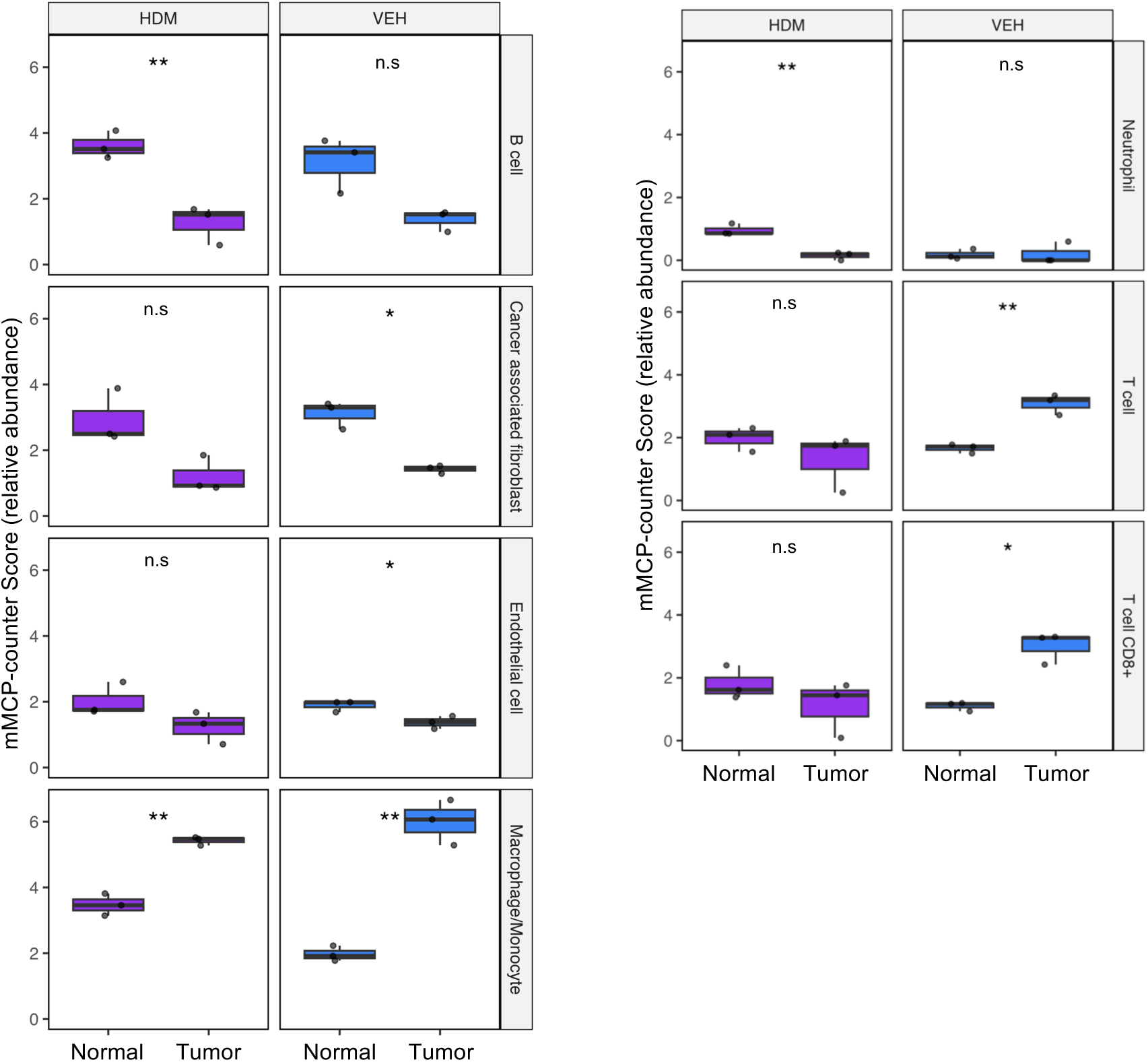
HDM Exposure Differentially Modulates Immune Cell Types in Tumors and Normal Adjacent Lung Tissues. Boxplots depict mMCP-counter scores for the indicated cell populations in tumors and normal lung tissues from mice treated with either HDM (purple) or VEH (blue). Boxes show the median (center line) and interquartile range (IQR); whiskers extend to 1.5× the IQR; points are individual samples (HDM, n = 3; VEH, n = 3). *p-*values were computed using a two-sided Welch’s *t*-test; n.s: non-significant, * *p* < 0.05, ** *p* < 0.01. Benjamini-Hochberg false discovery rate (FDR) was controlled at 10% (q ≤ 0.1) across cell types within each exposure, and panels were selected based on this FDR criterion.

**Figure S7.**
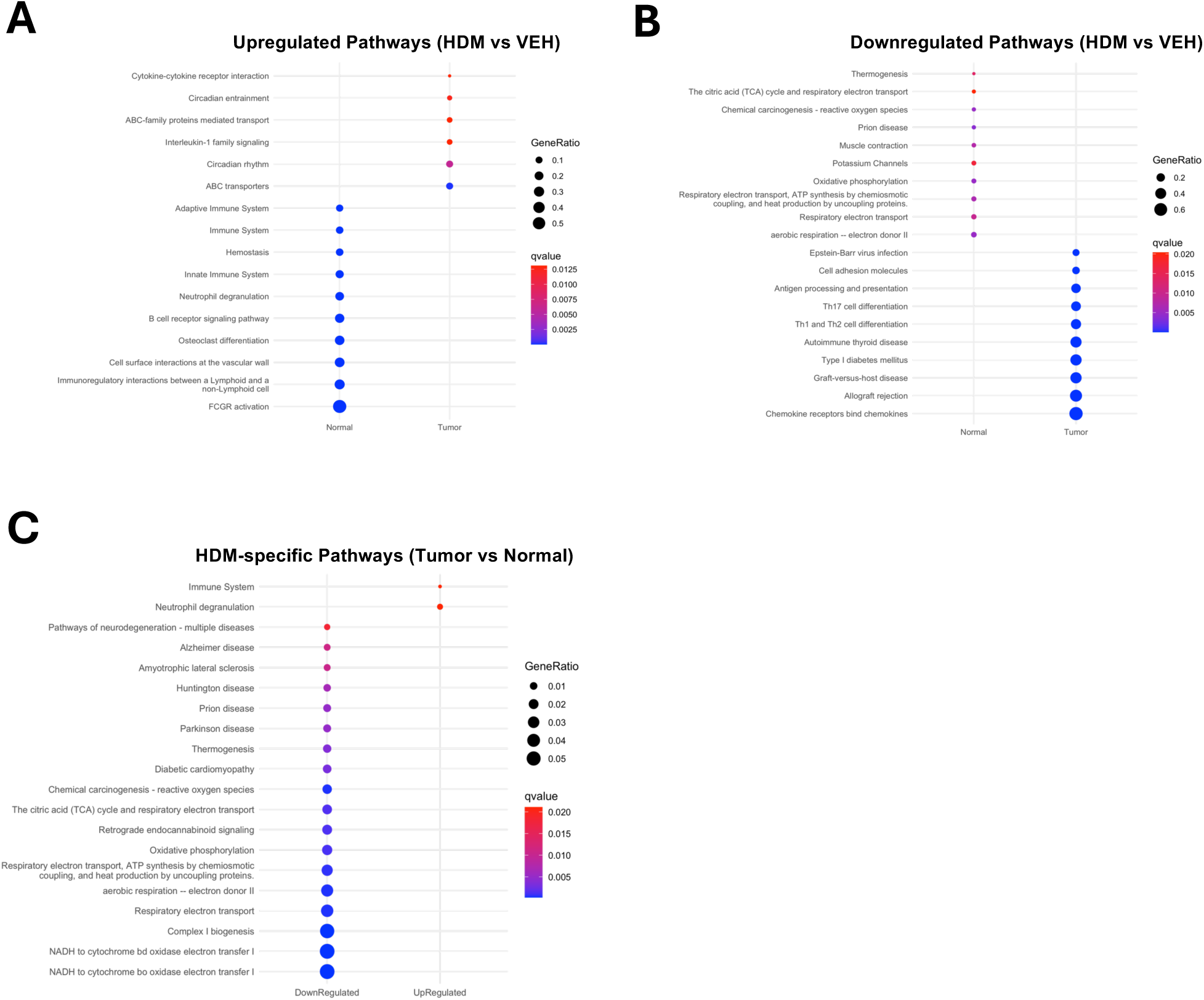
HDM Exposure Alters Signaling Pathways in Tumors and Normal Adjacent Lung Tissues. Pathway enrichment analysis of DEGs identified in tumor and normal adjacent lung tissues based on KEGG, Reactome, and MouseCyc pathway databases. The dot plots display all pathways (uncurated) that are either **A)** upregulated or **B)** downregulated in normal lung tissues or tumors following HDM exposure compared to VEH. **C)** Common pathways modulated by HDM in both normal lung tissues and tumors. Dot size represents the number of DEGs associated with each pathway, while dot color indicates statistical significance (q-value, calculated using the Benjamini-Hochberg false discovery rate).

**Figure S8.**
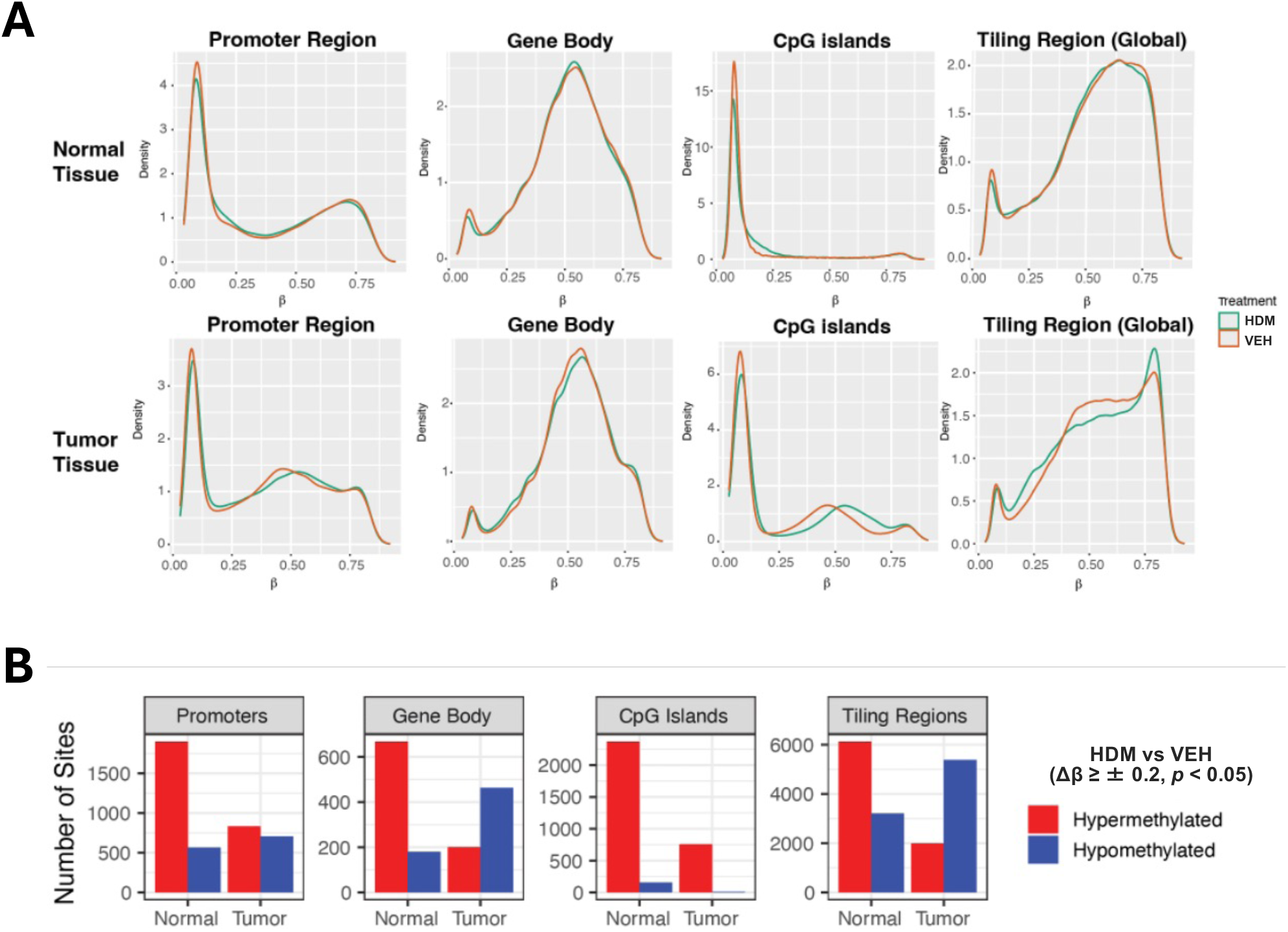
DNA methylation profiles of tumors and normal adjacent tissues exposed to HDM or VEH. **A)** β-value density plots. Kernel density curves showing the distribution of CpG β- values (0-1) across promoters, gene bodies, CpG islands, and genome-wide tiling regions for normal lung tissue (top row) and tumors (bottom row) under HDM (purple) and VEH (blue) conditions. **B)** Counts of differentially methylated sites (DMSs). Bar charts show the number of hypermethylated (red) and hypomethylated (blue) CpGs in each feature/tissue, defined as |Δβ| ≥ 0.20 and nominal *p* < 0.05 (Δβ = mean[HDM] − mean[VEH]).

